# A dose-dependent switch in translation capacity controls the transcription factor response to H_2_O_2_ stress

**DOI:** 10.64898/2026.01.28.701883

**Authors:** Chance Parkinson, Woody March-Steinman, Elizabeth Jose, Lisa Shanks, Andrew L. Paek

## Abstract

Hydrogen Peroxide (H_2_O_2_) stress activates transcription factors (TFs) in a dose-dependent manner, with distinct TFs activated in response to low versus high H_2_O_2_. Here, we show that high H₂O₂ imposes a translational constraint that prevents accumulation of TFs requiring *de novo* protein synthesis. Under low H_2_O_2_ conditions, TFs including p53, NRF2, and ATF4, accumulate and drive stress-responsive gene expression. In contrast, high H_2_O_2_ induces coordinated inhibition of translation initiation and elongation through activation of the integrated stress response (ISR), suppression of mTORC1 signaling, and activation of eEF2K, thereby blocking accumulation of these TFs. Inhibition of translation and repression of p53, NRF2, and ATF4 coincides with nuclear shuttling of pre-existing TFs, including FOXO1, NFAT1, and NF-κB. We propose that shuttling TFs provide a backup mechanism to respond to severe oxidative stress while translation is inhibited. Together, these findings identify translational control as a central switch governing transcription factor response to H_2_O_2_ stress.

## INTRODUCTION

Hydrogen peroxide (H_2_O_2_) is a reactive oxygen species (ROS) with well-established roles as a second messenger in processes such as wound healing, proliferation, and development^1–3^. However, dysregulated H_2_O_2_ production and the resulting redox imbalance can lead to DNA damage and oxidation of lipids and proteins which can potentially lead to cell death^4–6^. Oxidative damage has been linked to conditions such as chronic inflammation, ischemia-reperfusion injury, and oncogenesis, underscoring the necessity to tightly regulate H_2_O_2_ levels^7^.

Essential to the metazoan response to H_2_O_2_ stress is the activation of several transcription factors (TFs) including p53, NRF2, FOXO1 and NF-κB^8^. These TFs upregulate cytoprotective genes involved in NADPH/GSH production, cell cycle arrest, DNA damage repair, and autophagy. Each of these provides a critical step in alleviating oxidative stress. Our recent work elucidated a temporal and dose-dependent activation scheme for these and other TFs in response to acute H_2_O_2_ stress^9^. Low levels of H_2_O_2_ leads to rapid activation of p53, NRF2, JUN and ATF4. However, in response to higher levels of H_2_O_2_ stress, activation of these TFs is delayed while FOXO1, NF-κB, and NFAT1 are activated. These observations suggest that cells possess the remarkable ability to sense H₂O₂ concentrations and tailor their transcriptional response accordingly. Yet the mechanism for how this is achieved, and the cellular consequences remain unclear.

The tumor suppressor p53 is critical in the defense against genotoxic agents including H_2_O_2_ that can potentially cause deleterious mutations. In response to DNA damage, p53 activation induces various processes including cell cycle arrest, senescence, or apoptosis when damage is high^10^. Activation of p53 in response to H_2_O_2_ occurs through multiple DNA damage-dependent and independent mechanisms. For one, H_2_O_2_ can oxidize critical cysteine residues within apoptosis signal-regulated kinase-1 (Ask1), which occurs independent of DNA damage and stimulates p53 activation through activation of the stress-activated protein kinases (SAPKs) c-Jun N-terminal kinase (JNK) and p38^11–13^. Phosphorylation of p53 by JNK and p38 disrupts MDM2-dependent poly-ubiquitinylation of p53 and subsequent proteasomal degradation. The block in p53 degradation results in accumulation of p53 and an increase in transcriptional activity^14,15^.

Ataxia-telangiectasia mutated (ATM) protein kinase also has a well-established role in activating p53 in response to DNA double strand breaks. After being recruited to the Mre11-Rad50-Nbs1 (MRN) DNA repair complex, ATM phosphorylates p53 directly and additionally activates CHK2, which in turn phosphorylates p53^16^. Both phosphorylation events lead to stabilization of p53 by disrupting the interaction of p53 and MDM2. In addition, ATM can phosphorylate MDM2 at multiple sites further blocking binding to p53 and increasing MDM2 autoubiquitination and proteasomal degradation^17^. ATM is also activated through the formation of an H_2_O_2_-induced disulfide-cross-linked ATM dimer at cysteine 2991, independent of DNA damage or the MRN complex^18^. With multiple pathways stimulating p53 activation in response to H_2_O_2_, our previous finding that high levels of H_2_O_2_ stress suppress p53 accumulation, despite high levels of DNA damage, are striking^9^.

While H_2_O_2_ stress leads to the activation of the aforementioned TFs, cells also implement mechanisms to conserve energy by rapidly inhibiting the energetically costly process of translation^19^. This allows cells to reprogram gene expression and use available ATP for vital repair processes. In response to H_2_O_2_ stress, cytosolic ATP levels have been shown to drop to ∼50-80% of normal levels highlighting the need to limit unnecessary ATP usage and shunt energy towards antioxidant and repair pathways^20,21^. In both yeast and mammalian cells, several pathways are enacted to limit cap-dependent translation initiation and elongation including the integrated stress response (ISR) and eEF2K signaling^22,23^. Indeed, both yeast and mammalian cells activate these pathways causing global translation attenuation in response to H_2_O_2_ stress^24,25^. Additionally, mTORC1 signaling plays key roles in translational control and is inhibited in the face of many stressors including H_2_O_2_^26,27^. However, it is not known how cells coordinate translational control and TF activity in response to H_2_O_2_ stress as many transcription factors, including p53 require *de novo* translation in response to stress. Deciphering this relationship is key to understanding how cells counteract oxidative stress and may offer insights into diseases characterized by redox imbalance.

Here, we show that high H_2_O_2_ levels attenuate translation, which in turn blocks the accumulation of p53, NRF2, and ATF4. Focusing on p53, we find that its failure to accumulate at high H₂O₂ levels is not due to impaired transcription or enhanced degradation via autophagy or the proteasome. Live-cell imaging experiments further support the conclusion that a reduction in protein synthesis through translation, and not an increase in protein degradation, is responsible for the reduction of p53 at high levels of H_2_O_2_ stress. Indeed, activation of the ISR, inhibition of mTORC1 signaling, and activation of eEF2K are coordinated with suppression of p53, NRF2, and ATF4 at high doses of H_2_O_2_. The block in accumulation of these TFs coincides with shuttling of FOXO1, NF-KB, and NFAT1 to the nucleus. Strikingly, all three of the TFs activated rapidly under high H_2_O_2_ stress do not require de novo translation for their activation, as they are predominantly regulated by nuclear-to-cytoplasmic shuttling. The switch from p53, NRF2, and ATF4 under low H_2_O_2_ stress, to FOXO1, NF-KB, and NFAT1 under high H_2_O_2_ stress is accompanied by large changes in which target genes are activated. However, there is large overlap in the pathways activated by each group of TFs.

## RESULTS

### Suppression of p53 at high H_2_O_2_ concentrations is independent of MDM2 and the proteasome

To uncover the mechanism of p53 suppression at high levels of H_2_O_2_, we first tested the role of MDM2. We reasoned that though several stress-activated kinases, including ATM, p38 and JNK are known to stabilize p53 in response to H_2_O_2_, they do so by blocking MDM2-mediated proteasomal degradation of p53. Thus, if suppression of p53 occurs by inhibition of one or more upstream kinases, then p53 accumulation should be restored by adding Nutlin-3a, an MDM2 inhibitor.

We exposed cells to an H_2_O_2_ dose-response (0-200 μM), with or without a 10-minute pretreatment of Nutlin-3a and measured nuclear p53 levels by immunofluorescence (IF). As shown in a previous study, p53 activation exhibited an “inverted U” shaped dose-response to H_2_O_2_; low doses of H_2_O_2_ (40-100 μM) increased p53 protein levels, while high concentrations (150-200 μM) showed little to no p53 activation **(Fig. 1a**, left). Likewise, Nutlin-3a treatment increased p53 protein levels both in the absence of H_2_O_2_, and in response to low H_2_O_2_ concentrations (20-100μM, **Fig 1a**, right). In contrast, under high doses of H_2_O_2_ (150-200 μM) Nutlin-3A treatment did not result in an increase in p53 levels (**Fig. 1a**). Together these data suggest that MDM2 activity is not required to keep p53 levels low when cells are exposed to high H_2_O_2_.

**Figure 1.**
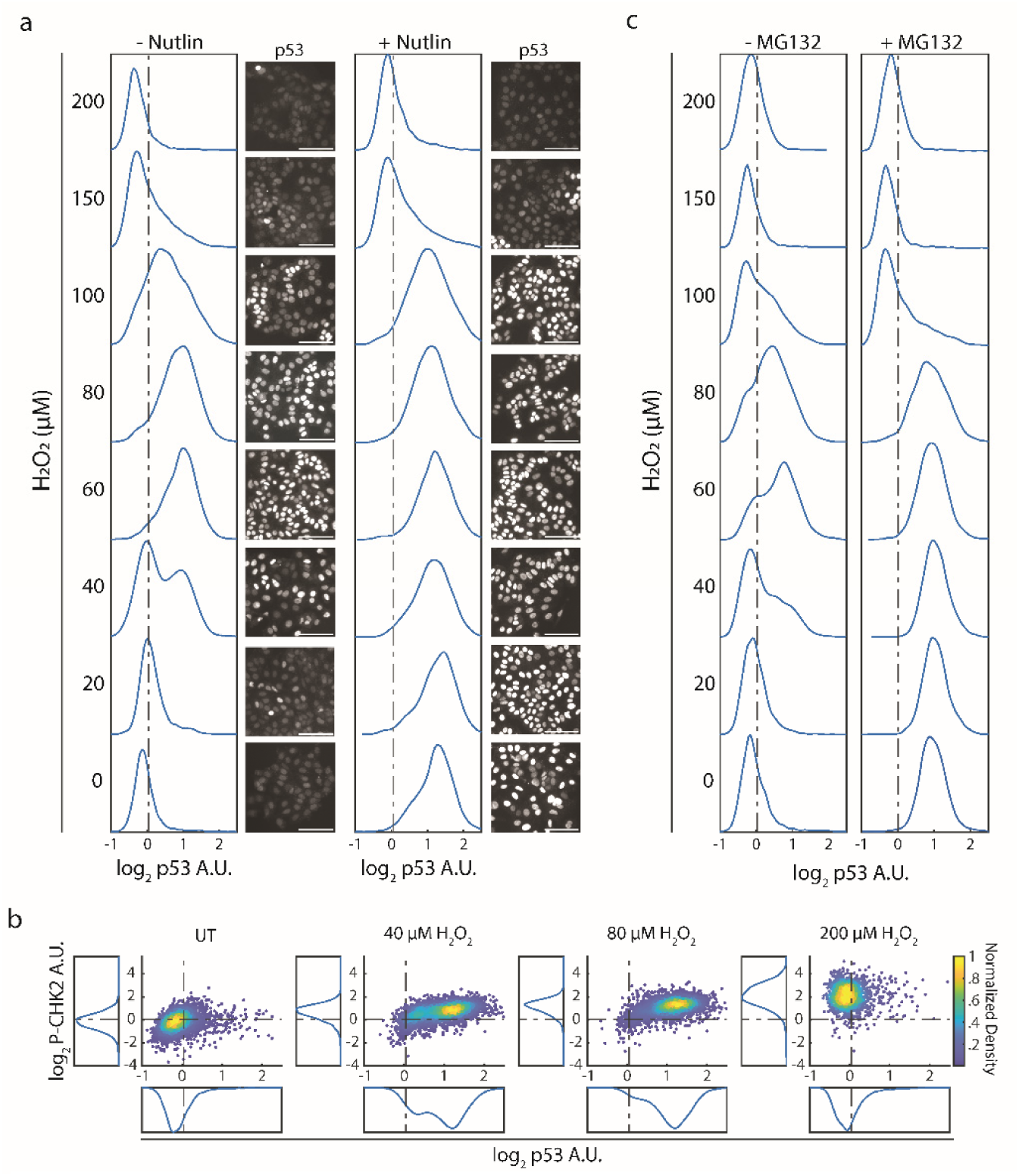
High H_2_O_2_ blocks p53 accumulation independent of the proteasome. (a) Population density plots of log2 nuclear p53 levels after treating MCF7 cells (n ≥ 3000) with the indicated H_2_O_2_ concentrations for 3hrs with (+) or without (-) Nutlin-3a (2.3 µM). IF staining of p53. Scale bar = 50 µm. (b) Density colored scatter plots and population density plots of IF data from MCF7 cells treated with the indicated concentrations of H2O2 for 3 hours. The log2 nuclear p53 levels (x-axis) and cellular P-CHK2 (T68) (y-axis) were measured. (c) As in (a) but treated with (+) or without (-) MG132 (5 µM). For all plots, values were normalized to the mean of the untreated samples (dashed line).

The fact that Nutlin-3a was unable to stabilize p53 protein in response to high H_2_O_2_ suggests that reduced activity of upstream kinases such as ATM is unlikely to be responsible for p53 suppression. In agreement with this, we observed an H_2_O_2_ dose-dependent increase in phospho-CHK2 (T68) levels, indicating an intact ATM signaling axis (**Fig. 1b**). To ensure that phospho-CHK2 is ATM dependent, we cotreated with the ATM inhibitor KU60019, which prevented phosphorylation of CHK2 across all H_2_O_2_ doses tested (**Supp Fig. 1a**). In addition, KU60019 diminished p53 accumulation at low doses of H_2_O_2_ (**Supp. Fig. 1b**), in agreement with previous studies that found multiple kinases (ATM, p38, JNK) promote p53 accumulation under H_2_O_2_ stress^12,13,18^. In contrast to phospho-CHK2 (T68), p53 levels were similar to untreated controls at the highest doses of H_2_O_2_ (200 μM) (**Fig. 1b**). This suggests that the block in p53 accumulation under high H_2_O_2_ occurs despite activation of ATM and CHK2.

We next tested whether suppression of p53 at high H_2_O_2_ is through proteasomal degradation, as p53 might be targeted by an alternative E3 ubiquitin ligase or through ubiquitin-independent proteasomal degradation^28^. Treating cells with the proteasome inhibitor, MG132, resulted in an increase in p53 protein levels similar to treatment with the MDM2 inhibitor Nutlin-3a (**Fig. 1c**). Yet combination treatment of high H_2_O_2_ (150-200 μM) with MG132 resulted in p53 levels comparable to untreated controls (**Fig. 1c**). This indicates that p53 suppression by high levels of H_2_O_2_ is independent of the proteasome.

In addition to p53, several other transcription factors, including ATF4 and NRF2, exhibit an “inverted U” shaped activation response to increasing levels of H_2_O_2_ (**Supp. Fig. 1c, d**). Both NRF2 and ATF4 are regulated in part by proteasomal degradation, thus we tested whether suppression of these transcription factors by high H_2_O_2_ is proteasome dependent^9,29^. Like p53, addition of the proteasome inhibitor MG132 alone resulted in accumulation of ATF4 and NRF2 yet failed to rescue ATF4 and NRF2 accumulation in response to high levels of H_2_O_2_ (**Supp. Fig. 1c, d**). Together these data show that inhibition of p53, ATF4 and NRF2 at high H_2_O_2_ is largely independent of the proteasome.

### Suppression of p53 at high H_2_O_2_ concentrations occurs independent of autophagy

We next reasoned that p53 could undergo degradation through autophagy at high levels of H_2_O_2_. To address this possibility, we used two chemical inhibitors of autophagy Chloroquine (CQ) and Bafilomycin A1 (BafA1). As above, we treated MCF7 cells with H_2_O_2_ in the presence of CQ or BafA1 and measured nuclear levels of p53. CQ treatment in the absence of H_2_O_2_ resulted in the formation of LC3 puncta indicative of immature autophagosomes and a block in autophagy (**Fig. 2a**)^30^. BafA1 treatment did not result in LC3 puncta as measured by IF (**Fig. 2a**). To confirm that BafA1 was blocking autophagy, we used the live-cell autophagic flux reporter, GFP-LC3-RFP-LC3ΔG^31^. GFP-LC3-RFP-LC3ΔG is cleaved by endogenous ATG4 proteases into equimolar amounts of GFP-LC3 and RFP-LC3ΔG. Once cleaved, GFP-LC3 can be degraded through the autophagosome while RFP-LC3ΔG cannot and serves as an internal control. Therefore, a decrease in the GFP/RFP ratio is indicative of an increase in autophagic flux. Using this probe, we observed that both BafA1 and CQ were able to block rapamycin- and H_2_O_2_-induced autophagy (**Fig. 2b**).

**Figure 2.**
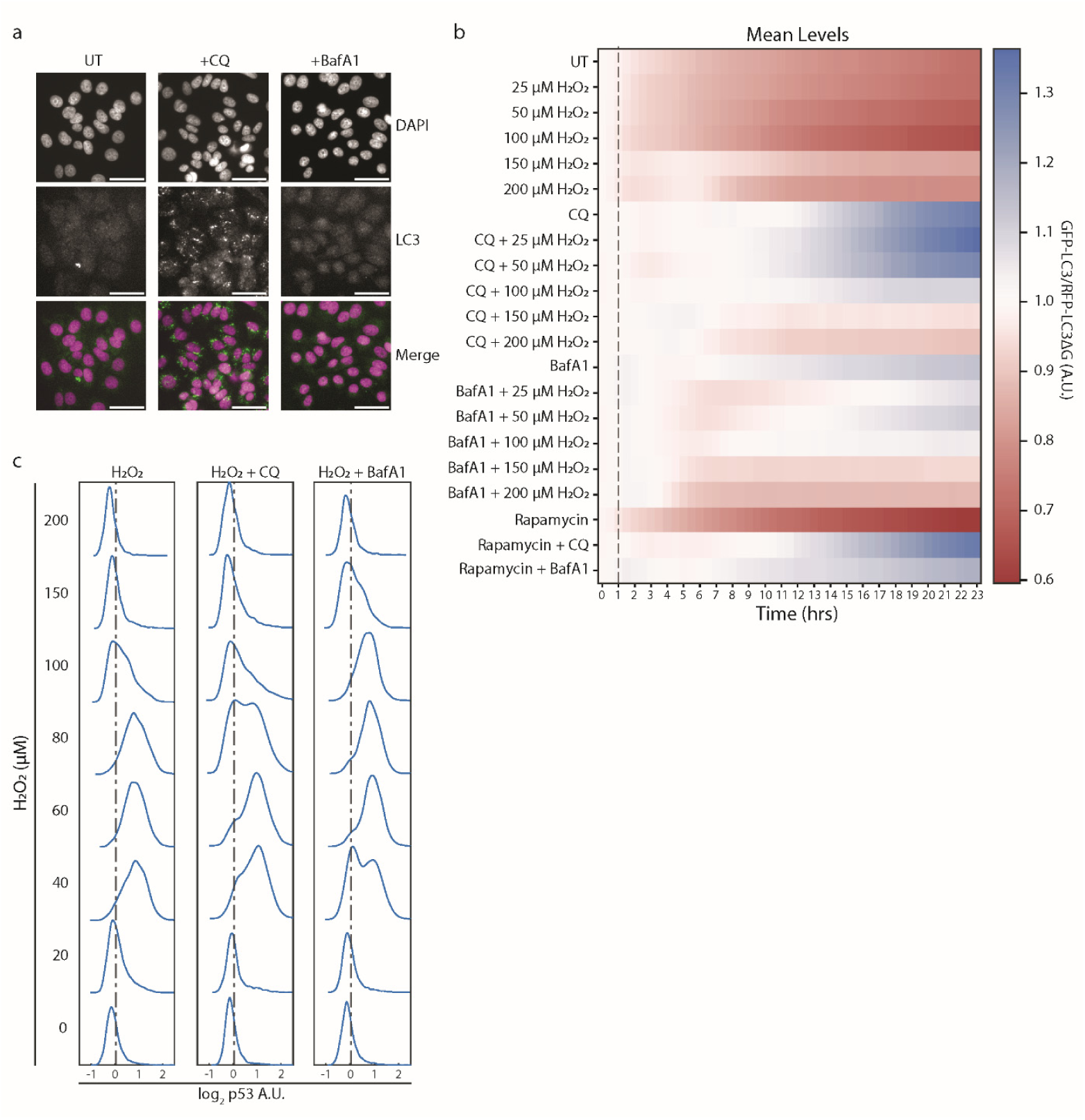
Autophagy is not required to block p53 accumulation at high H_2_O_2_. **(a)** IF staining of LC3 and DAPI after treating MCF7 cells with CQ (50 µM) or BafA1 (5 µM) for 1hr. Scale bar = 50 µm. **(b)** Heatmap generated from time-lapse microscopy data using the pMRX-IP-GFP-LC3-RFP-LC3ΔG reporter over 24hrs given the indicated treatment. The population mean of the GFP-LC3:RFP-LC3ΔG ratio was calculated at each time point (30min) and normalized to its starting value. Vertical gray dashed line represents H_2_O_2_ treatment time (t = 1hr) **(c)** Population density plots of log2 nuclear p53 levels when treated with the indicated H_2_O_2_ concentrations for 3hrs alone or after a 1hr pretreatment with CQ (50 µM) or BafA1 (5 µM). For (c), values were normalized to the mean of the untreated samples (dashed line).

We also observed that at higher doses of H_2_O_2_ (150-200 μM) there is a delay in the induction of autophagy for ∼5-7 hours marked by a GFP/RFP ratio close to 1 during this period (**Fig. 2b**). This suggests that p53 is an unlikely target for autophagic degradation during the initial response to high levels of H_2_O_2_. Indeed, despite effectively blocking autophagy, both CQ and BafA1 treatment did not lead to elevated p53 levels when combined with high doses of H_2_O_2_ (**Fig. 2c**). These data indicate that the suppression of p53 at high doses of H_2_O_2_ is independent of autophagy.

### Changes in transcription do not account for the suppression of p53 at high H_2_O_2_ concentrations

The above data indicates high H_2_O_2_ concentrations suppress p53 accumulation independent of proteasomal and autophagic degradation. Therefore, we hypothesized that changes in the transcription of the *TP53* gene could account for the dose-dependent response we observe. However, RNA sequencing data from our previous study shows minimal change in *TP53* transcript levels at all H_2_O_2_ concentrations tested^9^. To validate this, we utilized quantitative real-time PCR (qPCR) to measure changes in *TP53* mRNA levels at different H_2_O_2_ concentrations. Transcript levels of *TP53* increased modestly (1.2 – 1.4-fold) at doses of H_2_O_2_ that induce p53 accumulation. At the highest dose of H_2_O_2_ tested, *TP53* transcript levels were similar to untreated control cells, suggesting changes in *TP53* transcription are unlikely to explain the lack of p53 protein accumulation at high H_2_O_2_ doses (**Supp. Fig. 1e**). A similar trend was found when we analyzed transcript levels of *NFE2L2* and *ATF4*, the genes encoding NRF2 and ATF4, respectively (**Supp. Fig. 1f, g**). In contrast, transcript levels of the p53 target gene, *CDKN1A*, were dramatically affected by different concentrations of H_2_O_2_ with fold changes >20 at the doses where p53 is active **(Supp. Fig. 1h).** Finally, treatment of MCF7 cells with H_2_O_2_ in the presence of Actinomycin D, which intercalates into transcriptionally active regions of DNA and thereby abrogates RNA synthesis, did not prevent p53 accumulation at low to moderate H_2_O_2_ doses (20-100 μM) (**Supp. Fig. 1i**). Similar results have been reported for NRF2^32^. Together these data suggest that H_2_O_2_ regulates p53 post-transcriptionally, both its accumulation at low to moderate doses, and suppression at high doses.

### High doses of H_2_O_2_ block translation and prevent p53 production and degradation

Since neither degradation of p53 through the proteasome/autophagy nor a decrease in transcription of *TP53* mRNA is responsible for the suppression of p53 accumulation at high H_2_O_2_ doses, we reasoned that a decrease in p53 production through a block in translation is likely. To measure the dose-dependent impact of H_2_O_2_ on translation, we used the alkyne puromycin analog, O-propargyl-puromycin (OPP) linked to Alexa Fluor 647 Azide via click chemistry^33^.

We performed an H_2_O_2_ dose response and measured OPP incorporation into nascent peptides (a proxy for translation levels) and nuclear p53 levels in single cells by IF. We observed a dose-dependent decrease in translation upon treatment with H_2_O_2_ as shown in prior studies (**Fig. 3a, b**)^24^. At the highest dose of H_2_O_2_ tested (200 μM), OPP levels were ∼50% of control MCF7 cells. Treating cells with the translation inhibitor cycloheximide (CHX) decreased OPP fluorescence to ∼25% of control cells (**Fig. 3a, b**). These data suggest that H_2_O_2_ blocks some but not all translation in a dose-dependent manner. We observed a tight correlation between decreased translation and lack of p53 accumulation in single cells. At H_2_O_2_ doses above 60 μM there was a strong positive monotonic relationship between p53 protein levels and OPP incorporation (Spearman’s ρ = 0.62, p-value < 0.001; **Supp. Fig. 1k**). Additionally, IF images reveal that cells with low levels of p53 had low levels of OPP fluorescence and vice-versa (**Fig. 3c**). We further analyzed single-cell data by separating cells into 4 different bins based on OPP incorporation and p53 status (see **Supp. Fig. 1l** for activation thresholds). Less than 10% of cells had both high p53 levels and low OPP incorporation, indicating a strong reliance on translation for p53 accumulation in response to H_2_O_2_ (**Fig. 3a**). Indeed, pretreatment with CHX completely abrogated p53 accumulation in response to low and moderate levels of H_2_O_2_ (**Supp. Fig. 1j**).

**Figure 3.**
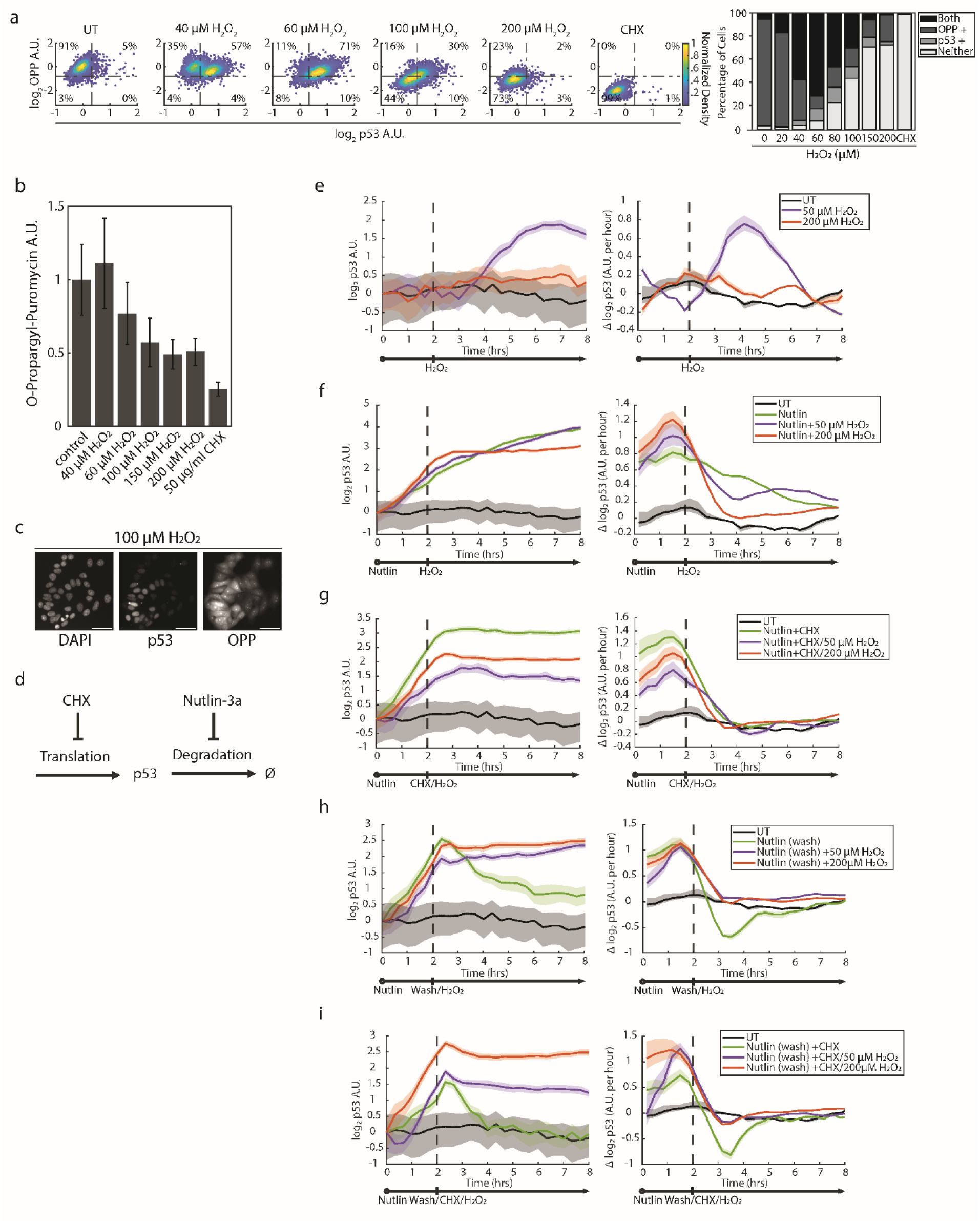
High H_2_O_2_ blocks p53 production and degradation 3) **(a)** Density colored scatter plots of the log2 nuclear p53 levels (x-axis) and log2 cellular OPP levels (y-axis) given the indicated treatment. Values were normalized to the mean of untreated controls. Activation thresholds (dashed lines) were determined using Otsu’s method on all single cell data as seen in Supp. Fig. 1k. On the right is a stacked bar graph which shows the percentage of cells containing high levels of both markers (“Both”), only one (e.g. – “p53+”), or neither (“Neither”) at the indicated treatment. **(b)** O-Propargyl-Puromycin (OPP) click chemistry assay showing median levels of AZDye647 cellular fluorescence after treatment with the indicated H_2_O_2_ concentrations for 3hrs. Values were normalized to the median of the control. Error bars = median absolute deviation. **(c)** IF images of cells treated with 100 µM H_2_O_2_ and stained with DAPI, p53 antibody, or AZDye647 (OPP). Scale bar = 50 µm. **(d)** Schematic illustrating how CHX and Nutlin-3a modulate p53 levels. CHX blocks new p53 translation and Nutlin-3a blocks p53 degradation. **(e-i)** Time-lapse microscopy traces of the p53-mScarlet-i3 reporter line over 8hrs given the indicated treatment. Left graphs represent the log2 of the population mean of nuclear p53-mScarlet-i3 intensity values (A.U.) calculated at each time point (20min) and normalized to the starting value (solid colored line). Right graphs show the smoothed derivative traces of the left p53-mScarlet-i3 mean traces (solid colored line). The shaded colored regions in both left and right graphs represent the standard error of the mean. Treatment schemes are below each graph. Vertical gray dashed lines represent treatment times. Cells were washed with 1ml of PBS once when indicated. Nutlin (10 µM), CHX (50 µg/ml), or H_2_O_2_ (50 µM or 200 µM) were added when indicated.

The correlation between reduced translation and lack of p53 accumulation at high H_2_O_2_ suggests translation attenuation limits p53 induction. To directly test the role of p53 translation and degradation in response to low and high H_2_O_2_, we performed a series of live-cell imaging experiments using cycloheximide (CHX) to block translation and the MDM2 inhibitor Nutlin-3a to block p53 degradation (**Fig. 3d**). To track p53 levels, we incorporated two lentiviral reporters in MCF7 cells: H2B-CFP for tracking nuclei and p53-mScarlet-I3 to track p53 levels. The mScarlet-I3 fluorescent protein has a maturation time of ∼2 minutes and therefore is well suited to capture changes in p53 translation rates^34^.

We first treated cells with a bolus of H_2_O_2_ and measured p53 levels every 20 minutes for 8 hours. As shown previously, 50 μM H_2_O_2_ led to an increase in p53 levels (∼4-fold increase by 3-5 hours), in contrast p53 levels remained flat in response to 200 μM H_2_O_2_ (**Fig. 3e**)^9^. We then tested how H_2_O_2_ affects the rate of p53 accumulation when cells are pretreated with the MDM2 inhibitor Nutlin-3a. We reasoned that if high H_2_O_2_ blocks p53 translation, then it should halt the increase in p53 from Nutlin-3a treatment. Indeed, p53 levels stabilized ∼1 hour after treatment with 200 μM H_2_O_2_, remaining constant for the remainder of the experiment (**Fig. 3f**, left). This is reflected in the derivative of p53 levels (Δp53/hour), which rapidly approaches and remains near 0 after 200 µM H_2_O_2_ treatment, indicating p53 levels have reached a steady state (**Fig. 3f**, right). In contrast, cells treated with Nutlin-3a alone, or in combination with 50μM H_2_O_2_, displayed a continued increase in p53 levels, and a positive derivative, indicating continued translation of p53 (**Fig. 3f**). To compare these results to a complete shutdown of translation, we repeated this experiment but added cycloheximide in addition to H_2_O_2_ **(Fig. 3g).** Adding cycloheximide resulted in a similar stabilization of p53 levels in the absence or presence of H_2_O_2_ regardless of dose **(Fig. 3g)**. Indeed, the accumulation of p53 in response to 200 μM H_2_O_2_ was not affected by addition of cycloheximide, suggesting little to no translation of p53 at this dose **(Fig. 3f, g** compare red traces).

Finally, to ensure that MDM2 dependent degradation of p53 does not affect the response to high H_2_O_2_, we washed Nultin-3a out of the media prior to treatment with H_2_O_2_ or co-treatment with H_2_O_2_ and CHX. In the absence of H_2_O_2_, Nutlin-3a washout resulted in a decay of p53 signal over time, reflecting the rapid degradation of p53 by MDM2. In contrast, treatment with H_2_O_2_ after Nutlin-3a washout resulted in stabilization of p53, suggesting that both high and low doses of H_2_O_2_ block degradation of p53 **(Fig. 3h)**. Similar results occurred when CHX was added after Nutlin-3a washout (**Fig. 3i**). Together these data suggest that high H_2_O_2_ blocks both production and degradation of p53, and the lack of p53 accumulation in response to high H_2_O_2_ likely occurs by attenuating translation.

### High doses of H_2_O_2_ block translation and prevent production and degradation of NRF2 and ATF4

To test if NRF2 and ATF4 protein accumulation is also suppressed by translation inhibition at high levels of H_2_O_2_, we used immunofluorescence together with chemical inhibitors that stabilize these proteins by blocking their degradation. Specifically, we inhibited KEAP1, the E3 ligase that targets NRF2 for proteasomal degradation, or the proteasome to modulate NRF2 and ATF4 levels, respectively. Pretreatment with the KEAP1 inhibitor KI696 for two hours increased NRF2 levels across all H₂O₂ doses tested (**Supp. Fig. 2a**, middle). In contrast, treating with KI696 one hour after H₂O₂ treatment increased NRF2 levels for low doses of H_2_O_2_, but failed to increase NRF2 to the same levels when exposed to high H₂O₂ doses (**Supp. Fig. 2a**, right). Similarly, pretreatment with the proteasome inhibitor MG132 increased ATF4 levels at all doses H_2_O_2_ (**Supp. Fig. 2b**, middle), but only increased ATF4 levels for low doses of H₂O₂ when MG132 was added one hour after H₂O₂ (**Supp. Fig. 2b**, right). As expected, p53 was impacted by MG132 treatments in a similar fashion to ATF4 (**Supp. Fig. 2c**). Together, these data support the conclusion that high doses of H_2_O_2_ blunt accumulation of the stress responsive TFs NRF2, ATF4, and p53 by inhibiting translation rather than increasing protein degradation.

### H_2_O_2_-induced suppression of p53 coincides with the activation of the integrated stress response, the elongation inhibitor eEEF2K, and inhibition of mTORC1

We next measured how different translation control pathways correspond with p53 suppression under H_2_O_2_ stress. We began by assessing the integrated stress response (ISR). The ISR is characterized by four stress-responsive kinases (GCN2, PERK, PKR, and HRI) that phosphorylate the α subunit of eukaryotic initiation factor 2 (eIF2α) on serine 51 (S51)^35^. Phosphorylation of eIF2α-S51 prevents formation of the ternary complex, which is comprised of eIF2, the initiator Met-tRNA_i_^Met^, and GTP. The ternary complex is required for efficient cap-dependent translation initiation, and p53 translation is partially cap dependent^35,36^. Therefore, we examined if ternary complex formation was compromised at high concentrations of H_2_O_2_ by using IF to measure phosphorylation of eIF2α at S51 (P-eIF2α). We observed a dose-dependent increase in the levels of P-eIF2α when MCF7 cells were treated with H_2_O_2_. Furthermore, P-eIF2α and p53 accumulation were mutually exclusive as cells with high P-eIF2α had low levels of p53 and vice-versa (**Fig. 4a**).

**Figure 4.**
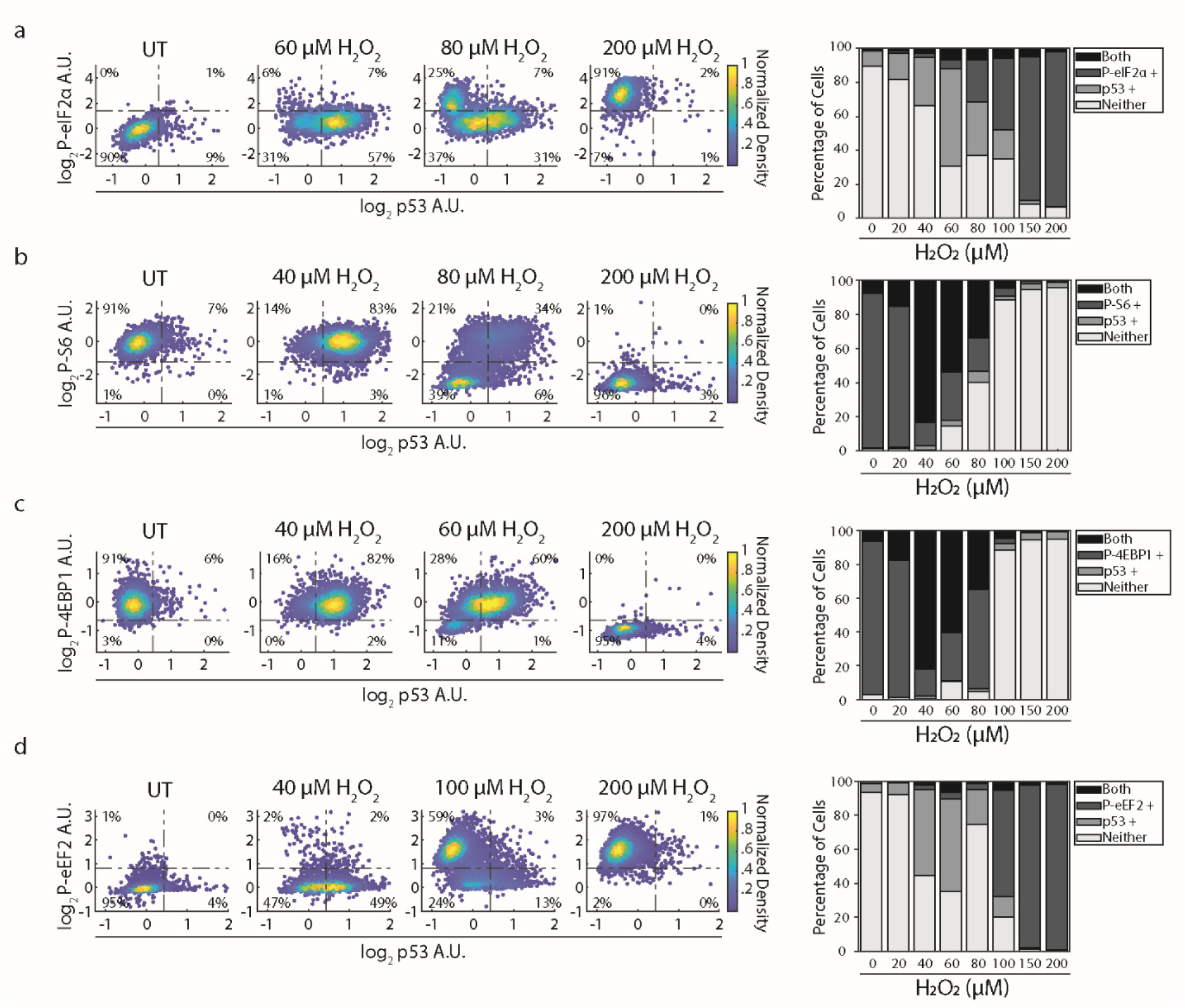
H_2_O_2_-induced suppression of p53 coincides with activation of multiple translation attenuation pathways. **(a-d)** MCF7 cells were treated with the indicated H_2_O_2_ concentrations for 3hrs. IF was used to measure **(a)** log2 nuclear p53 levels (x-axis) and the log2 cytoplasmic P-eIF2α (S51) (y-axis), **(b)** log2 nuclear p53 levels (x-axis) and the log2 cytoplasmic P-S6 (S256/236) (y-axis), **(c)** log2 nuclear p53 levels (x-axis) and the log2 cellular P-4EBP1 (T37/46), and **(d)** log2 nuclear p53 levels (x-axis) and the log2 cytoplasmic P-eEF2 (T56) (y-axis). For **(a-d)**, all values were normalized to the mean of untreated controls. Activation thresholds (dashed lines) were determined using Otsu’s method on all single cell data as seen in Supp. Fig. 1k. On the right are stacked bar graphs which show the percentage of cells containing high levels of both markers (“Both”), only one (e.g. – “p53+”), or neither (“Neither”) at the indicated concentrations of H_2_O_2_.

In addition to the ISR, the mammalian target of rapamycin complex 1 (mTORC1) controls cap-dependent translation initiation and is known to be regulated through oxidation-dependent mechanisms^26,27^. To determine the impact of H_2_O_2_ on mTORC1 activity, we measured mTORC1-dependent phosphorylation events which upregulate cap-dependent translation: phosphorylation of ribosomal protein S6 at Ser235/236 (P-S6) and phosphorylation of eukaryotic translation initiation factor 4E-binding protein 1 at Thr37/46 (P-4EBP1). We found that both P-S6 and P-4EBP1 levels decreased in a dose-dependent manner in response to H_2_O_2_, and suppression of p53 at high doses of H_2_O_2_ strongly coincided with the decrease in P-S6 and P-4EBP1; <6% of cells had both high levels of nuclear p53 and low levels of P-S6 or P-4EBP1 **(Fig. 4b, c).** These data show that multiple pathways (i.e. - ISR and mTORC1) which impact cap-dependent translation initiation are affected by high doses of H_2_O_2_ in a manner that coincides with translation attenuation and p53 suppression.

Others have noted a post-initiation translation block in response to H_2_O_2_, so we next tested if this was associated with p53 inhibition. The eukaryotic elongation factor 2 (eEF2) is a GTP-dependent protein critical for the transfer of the nascent peptide from the A to the P site of the ribosome^37^. eEF2 activity is inhibited by phosphorylation of Thr56 by the kinase eEF2K^38^. Therefore, we measured levels of phosphorylated eEF2 at Thr56 (P-eEF2) in response to H_2_O_2_ to determine if a translation elongation block occurs in response to high H_2_O_2_. Indeed, we observed a dose-dependent increase of P-eEF2 in response to H_2_O_2_. Like P-eIF2α, phosphorylation of eEF2 and activation of p53 were largely mutually exclusive in single cells **(Fig. 4d).**

In summary, multiple mechanisms of translational repression are engaged in response to high H_2_O_2_. This includes inhibition of cap dependent translation through the ISR and inhibition of mTORC1, and activation of the translation elongation inhibitor eEF2K.

### The correlation between translation repression mechanisms and the block in p53 accumulation is not limited to MCF7 cells

The observed pattern of ISR and eEF2K activation, mTORC1 inhibition, and p53 suppression was also observed in MCF10A and U2OS cells (**Supp. Fig. 3a-h**). In line with our previous study, it took much higher levels of H_2_O_2_ to activate translation repression pathways and inhibit p53 accumulation in U2OS cells^9^. It is unclear why U2OS cells are more resistant to H_2_O_2_ than MCF7 or MCF10A cells. However, these data show that the attenuation of translation initiation and elongation coincides with suppression of p53 accumulation at high doses of H_2_O_2_, and that this is likely a general response to severe H_2_O_2_ stress.

### Phosphorylation of eIF2α is sufficient to block H_2_O_2_ induced p53 accumulation but not necessary

We next tested whether phosphorylation of eIF2α is required to block accumulation of p53 at high H_2_O_2_. Using small molecule inhibitors, we found that phosphorylation of eIF2α under H_2_O_2_ stress is facilitated in part by PERK, but mostly GCN2 (**Fig. 5a**). A combination treatment of both PERK and GCN2 inhibitors completely blocked P-eIF2α in response to high H_2_O_2_, yet this failed to restore nuclear p53 levels at high H_2_O_2_ (**Fig. 5a** & **Supp. Fig. 4a**). Moreover, translation was still attenuated by H_2_O_2_ in cells treated with the PERK and GCN2 inhibitors as measured by the OPP incorporation assay (**Supp. Fig. 4b)**.

**Figure 5.**
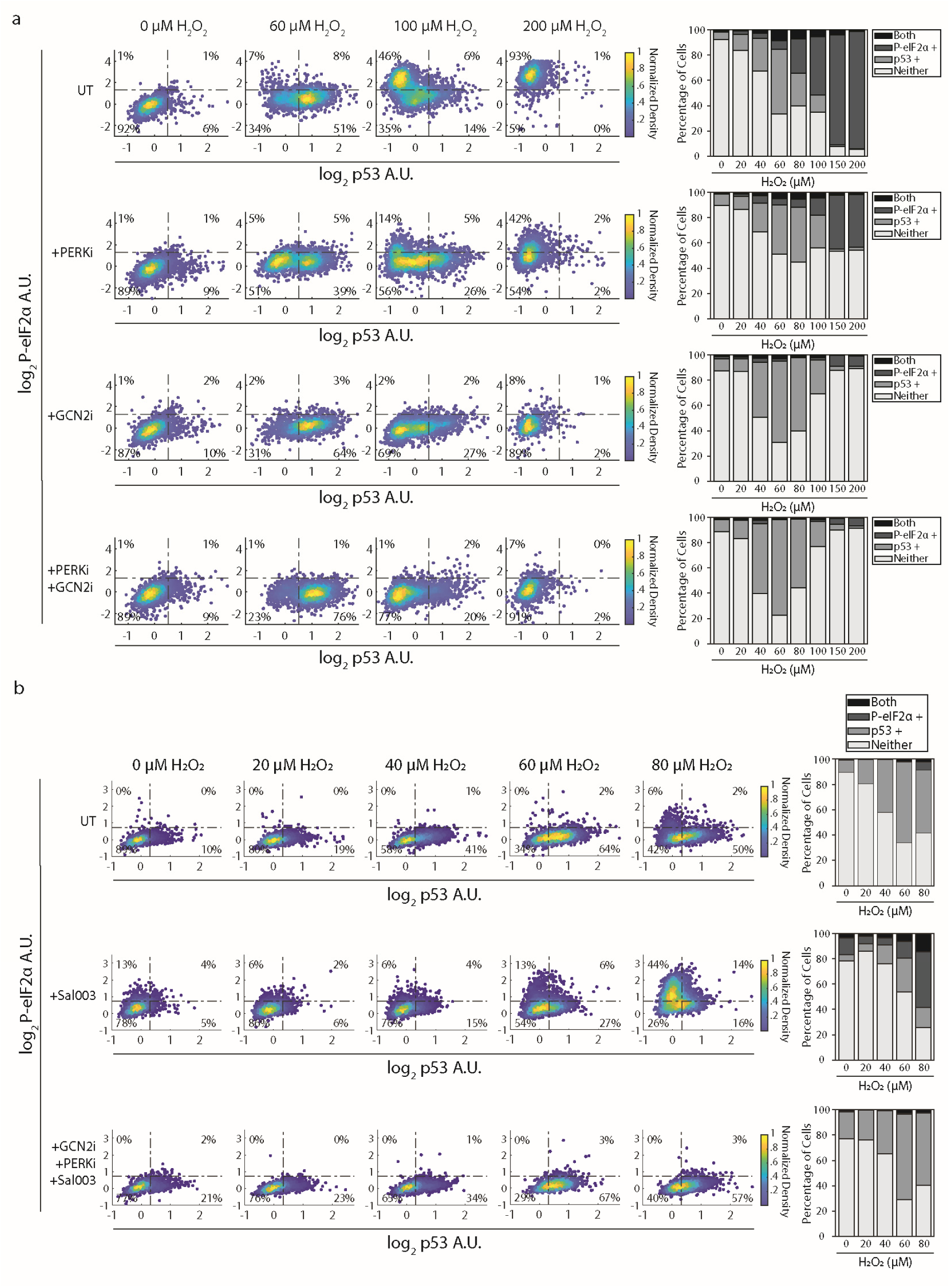
The ISR is sufficient but not required for blocking p53 accumulation. **(a)** MCF7 cells were treated with the indicated H_2_O_2_ concentrations for 3hrs with or without PERK and/or GCN2 inhibitors (+PERKi 1.25 µM; +GCN2i 5 µM). Density colored scattered plots from IF data measuring log2 nuclear p53 levels (x-axis) and log2 cytoplasmic P-eIF2α (S51) (y-axis) at each of the specified H_2_O_2_ concentrations. **(b)** MCF7 cells were first treated with or without PERK and GCN2 inhibitors (+PERKi 1.25 µM; +GCN2i 5 µM) for 10min followed by Sal003 (50 µM) treatment for 1hr. Cells were then treated with H_2_O_2_ for 3 hrs. Density colored scattered plots from IF data measuring log2 nuclear p53 levels (x-axis) and log2 cytoplasmic P-eIF2α (S51) (y-axis) at each of the specified H_2_O_2_ concentrations. On the right of (a, b) are stacked bar graphs which show the percentage of cells containing high levels of both markers (“Both”), only one (e.g. – “p53+”), or neither (“Neither”) at the indicated concentrations of H2O2. For (a, b), activation thresholds (dashed lines) were determined using Otsu’s method on all single cell data as seen in Supp. Fig. 1k.

The fact that P-eIF2α dependent inhibition of cap-dependent translation is not required to block p53 accumulation is perhaps not surprising as multiple mechanisms of translational repression are engaged in response to high H_2_O_2_. Thus, we sought to test whether activation of the ISR is sufficient for blocking p53 accumulation. To do this we first utilized the translation inhibitor arsenite, a well-known inducer of P-eIF2α. Pretreatment with arsenite followed by H_2_O_2_ blocked p53 accumulation in the presence of any H_2_O_2_ dose tested (**Supp. Fig. 4c**). However, blocking P-eIF2α with a GCN2 inhibitor, failed to restore p53 accumulation in response to arsenite (**Supp. Fig. 4c**). This is likely because arsenite induces stress granules and sequesters other vital translation initiation components which could be blocking translation of p53 mRNA. It should be noted that we did not observe G3BP-positive stress granules at any of the H_2_O_2_ concentrations tested in this study. This is in direct contrast to arsenite treated cells (**Supp. Fig. 4d**).

We next tested whether the eIF2α phosphatase inhibitor Sal003 was sufficient to induce P-eIF2α and block H_2_O_2_ induced p53 accumulation^39^. Cells were treated with Sal003 for one hour followed by an H_2_O_2_ dose response. Sal003 alone induced a modest increase of P-eIF2α levels which was exacerbated by low doses of H_2_O_2_ (**Fig. 5b**). The increase in P-eIF2α with Sal003 was associated with a reduction in p53 accumulation. Pre-treatment with GCN2 and PERK inhibitors prevented the Sal003-induced increase in P-eIF2α and restored p53 to H_2_O_2_-only levels (**Fig. 5b**). Thus, phosphorylation of eIF2α through PERK and/or GCN2 is sufficient to prevent p53 accumulation in response H_2_O_2_ stress but is not required to block p53 accumulation under high levels of H_2_O_2_.

### mTORC1 inhibition is not sufficient to block p53 accumulation at low doses of H_2_O_2_

The Tuberous sclerosis complex 2, or Tuberin (TSC2), is a negative regulator of mTORC1 signaling^40^. TSC2 knockout cell lines maintain mTORC1 signaling even in the absence of serum^41^. Thus, we first tested if TSC2 knockouts maintain mTORC1 signaling under high level of H_2_O_2_ stress. However, we found that a CRISPR-Cas9 mediated knockout of TSC2 failed to restore P-S6 and P-4EBP1 in response to high levels of H_2_O_2_ (**Supp. Fig. 5a, b**). P53 accumulation was also suppressed at high H_2_O_2_ doses in TSC2 knockout and parental MCF7 cells (**Supp. Fig. 5b**). Therefore, we next tested whether knockout of 4EBP1 could alleviate p53 suppression under high H_2_O_2_ stress. 4EBP1 represses cap-dependent translation by binding to the eukaryotic translation initiation factor 4E (eIF4E). mTORC1 phosphorylation of 4EBP1 releases eIF4E from 4EBP1, promoting cap-dependent translation^42^. Using CRISPR-Cas9, we generated a 4EBP1 knockout cell line and found no change in p53 levels across the H_2_O_2_ dose response **(Supp. Fig. 5a, c**). Additionally, neither the TSC2 knockout nor 4EBP1 knockout were able to restore translation as measured by the OPP assay (**Supp. Fig. 5b, c).** Therefore, 4EBP1 is not required to block p53 translation at high levels of H_2_O_2_.

To test whether inhibiting mTOR activity is sufficient to block p53 accumulation at low doses of H_2_O_2_ stress, we utilized the small molecules rapamycin (mTORC1 specific inhibitor) and AZD8055 (mTORC1 and mTORC2 specific inhibitor). Pretreatment with rapamycin resulted in a decrease of P-S6 but not P-4EBP1 as shown previously (**Supp. Fig. 5d, e**)^43^. When cells were pretreated with rapamycin and then treated with H_2_O_2_, p53 still accumulated to levels similar to H_2_O_2_ treated cells in the absence of rapamycin (**Fig. 6a**). Treatment with AZD8055 blocks both mTORC1-dependent phosphorylation of S6 and 4EBP1 (**Supp. Fig. 5d, e**). However, pretreatment with AZD8055 did not abolish p53 accumulation in the presence of low levels of H_2_O_2_ stress (**Fig. 6b**). These data indicate inhibition of mTORC1 activity alone is not sufficient to block p53 translation in the presence of low to moderate H_2_O_2_ stress.

**Figure 6.**
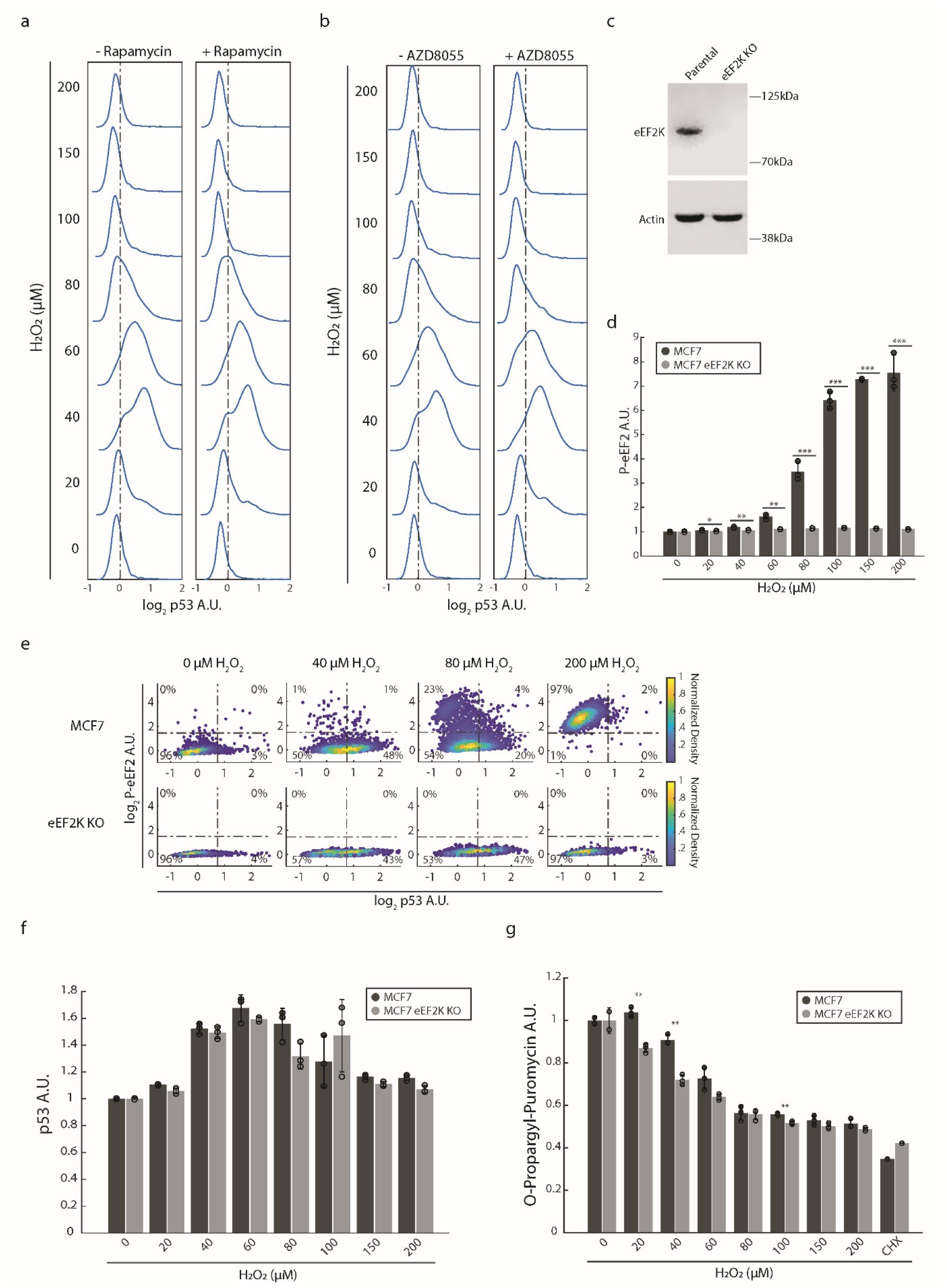
Inhibition of mTORC1 is not sufficient and eEF2K is not required to block p53 accumulation. **(a)** Population density plots of log2 nuclear p53 levels after treating MCF7 cells with the indicated H_2_O_2_ concentrations for 3hrs with (+) or without (-) Rapamycin (220nM). **(b)** As in (a) but treated with (+) or without (-) AZD8055 (100nM). **(c)** Western blot stained for eEF2K and actin in MCF7 parental cells or a clonal line containing an eEF2K CRISPR-Cas9 mediated knockout. **(d)** Bar graph showing normalized cytoplasmic levels of P-eEF2 (T56) for the indicated cell lines treated with H_2_O_2_. **(e)** Density colored scatter plots of the log2 nuclear p53 levels (x-axis) and the log2 cytoplasmic P-eEF2 (T56) levels (y-axis) for the indicated cell line treated with H_2_O_2_ for 3hrs. Activation thresholds (dashed lines) were determined using Otsu’s method on all single cell data as seen in Supp. Fig. 1k. **(f)** Bar graph showing normalized nuclear levels of p53 for the indicated cell lines treated with H_2_O_2_ for 3hrs. **(g)** Bar graph showing normalized cellular levels of OPP for the indicated cell lines treated with H_2_O_2_ for 3hrs or CHX (50 µg/ml) for 30min. For bar graphs (d, f, g), values were normalized to the untreated condition for each cell line. Error bars represent the sample standard deviation. Dots represent replicate means. P-values were acquired by first performing a two-sample t-test then adjusted for multiple hypotheses using the Benjamini-Hochberg procedure (* ≤ 0.05, ** ≤ 0.01, *** ≤ 0.001).

### eEF2 phosphorylation is not necessary to prevent p53 accumulation in response to high H_2_O_2_ levels

We next tested whether P-eEF2 dependent inhibition of translation elongation is required for the block in p53 accumulation at high H_2_O_2_. We used CRISPR-Cas9 to knock out the gene encoding eEF2K; the kinase that phosphorylates eEF2 at T56 (**Fig. 6c**). Knocking out eEF2K prevented eEF2 phosphorylation in response to H_2_O_2_ (**Fig. 6d)**. Using this knockout line, we did not observe a significant increase in nuclear p53 levels at moderate to high H_2_O_2_ doses when compared to the parental MCF7 cell line (**Fig. 6e, f**). Additionally, preventing phosphorylation of eEF2 did not restore translation as measured by the OPP incorporation assay (**Fig. 6g**). Therefore, we conclude that eEF2 phosphorylation is not necessary to block p53 accumulation or translation at high levels of H_2_O_2_. This is in contrast to another group’s findings which showed that an eEF2K KO in MEFs and fission yeast was sufficient to restore translation^25^. It is possible that human cells rely more heavily on other translation control pathways (e.g. – ISR, mTORC1, or others) than mice and yeast, but this requires further investigation.

### Drugs that induce phosphorylation of eEF2 block p53 accumulation at low doses of H_2_O_2_ in parental and eEF2K knockout cells

We next tested whether induction of P-eEF2 is sufficient to block p53 accumulation at low levels of H_2_O_2_ stress. Using the eEF2K activator Nelfinavir (NFR), we found that induction of P-eEF2 is able to suppress p53 translation at low levels of H_2_O_2_ (**Supp. Fig. 6a**)^44^. However, this suppression also occurred in the eEF2K knockout cell line indicating this finding is not the result of eEF2K activation specifically (**Supp. Fig. 6a**). We next turned to another drug, NH125, reported to activate eEF2K-dependent phosphorylation of eEF2 at high concentrations^45^. Indeed, we observed NH125 induces moderate activation of eEF2K and thus phosphorylation of eEF2 (**Supp. Fig. 6b**). Pretreatment with NH125 and then H_2_O_2_ treatment resulted in a block of p53 accumulation at all doses of H_2_O_2_ in both parental and eEF2K knockout cells as seen above with NFR (**Supp. Fig. 6b)**. Therefore, we are unable to state definitively whether phosphorylation of eEF2 is sufficient to block accumulation of p53. Additionally, researchers should use caution when utilizing these drugs to activate eEF2K signaling as off-target effects are likely.

### Cytoplasmic-to-nuclear shuttling of several TFs coincides with translation attenuation

In a prior study, we identified two TF modules that respond to H₂O₂ in a dose and time-dependent manner^9^. At low H₂O₂, p53, NRF2, JUN, and ATF4 (the “p53 group”) are rapidly activated, whereas FOXO1, NF-κB, and NFAT1 (the “FOXO1 group”) remain inactive. In contrast, high H₂O₂ initially represses the p53 group while activating the FOXO1 group via cytoplasmic-to-nuclear translocation. The duration of FOXO1 group nuclear accumulation increases with H₂O₂ dose, and once the FOXO1 group exits the nucleus, the p53 group of TFs begin to accumulate. Our observation that repression of the p53 group is driven by translational repression provides a mechanistic basis for the separation of these two TF responses. All p53 group TFs require *de novo* translation for activation, either through stabilization by inhibition of protein degradation (p53, NRF2, ATF4)^29,46,47^ or increased mRNA transcription and/or translation (NRF2, ATF4, JUN)^48–50^. In contrast, the FOXO1 group of TFs are activated by nuclear shuttling and do not require new protein synthesis. Indeed, all markers of translation inhibition examined here, including activation of the ISR and eEF2K, and inhibition of mTORC1 signaling, coincide with nuclear shuttling of FOXO1 (**Fig. 7a-d**).

**Figure 7.**
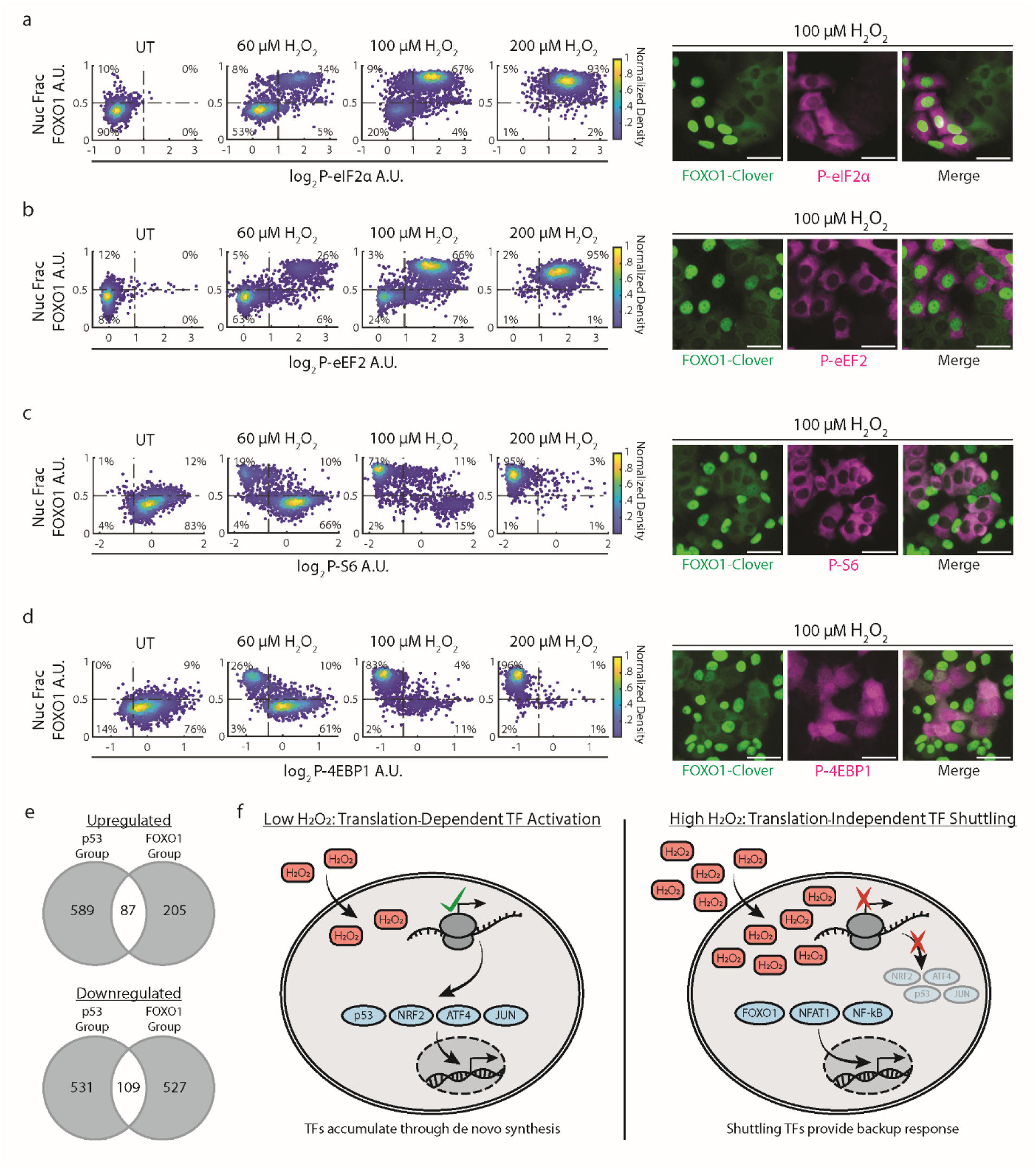
Translation attenuation coincides with shuttling of FOXO1. Data was generated from MCF7 cells expressing a lentiviral reporter of FOXO1-Clover **(a)** Left: density colored scatter plots of the nuclear fraction (“Nuc Frac”) of FOXO1-Clover (x-axis) and log2 cytoplasmic P-eIF2α (S51) levels (y-axis) after treatment with H_2_O_2_ for 3hrs. Right: IF images of cells treated with 100 µM H_2_O_2_ and probed for FOXO1-Clover and P-eIF2α (S51). **(b)** Same as in (a) but measuring the log2 cytoplasmic P-eEF2 (T56) (y-axis). **(c)** Same as in (a) but measuring the log2 cytoplasmic P-S6 (S256/236) (y-axis). **(c)** Same as in (a) but measuring the log2 cellular P-4EBP1 (T37/46) (y-axis). For (a-d), activation thresholds (dashed lines) for all but Nuc Frac FOXO1 were determined using Otsu’s method on all single cell data as seen in Supp. Fig. 1k. Nuc Frac FOXO1 activation threshold was set at 0.5. Scale bar = 50 µm. **(e)** Venn diagram of the number of significantly up- and downregulated genes (p_adj_ < 1e-5, |Log2FoldChange| > 0.2) for both the “p53 Group” and “FOXO1 Group.” RNAseq data for (e) was acquired in our recent work (Jose et al, 2024). “p53 Group” represents data taken from cells with p53 accumulation after H_2_O_2_ treatment. “FOXO1 Group” represents data taken from cells without p53 accumulation after H_2_O_2_ treatment; these cells instead activate TFs such as FOXO1, NFAT1, and NF-κB. **(f)** Model showing activation of TFs that rely on *de novo* synthesis at low concentrations of H_2_O_2_ (left), while high doses of H_2_O_2_ inhibit translation and therefore rely on cytoplasmic-to-nuclear shuttling of TFs to transcribe stress-response genes (right).

The coordinated attenuation in translation, repression of the p53 group of TFs, and nuclear translocation of the FOXO1 group likely serves multiple purposes. First, activation of the FOXO1 group may act as a back-up mechanism to drive critical gene expression changes when the p53 group is no longer translated. Second, the FOXO1 group may regulate a distinct transcriptional program specifically required during severe H_2_O_2_ stress. To explore these two models, we analyzed our previously published RNA-seq data^9^. The data set took advantage of our finding that activation of the two TF groups is controlled by the peroxiredoxin (PRDX) and sulfiredoxin (SRXN1) proteins. High H_2_O_2_ leads to hyperoxidation and inactivation of PRDX1 and PRDX2 proteins, which in turn results in repression of the p53 group of TFs, and activation of the FOXO1 group. PRDX1/2 hyperoxidation is reversed by the sulfiredoxin (SRXN1) enzyme. Thus, using the same dose of H_2_O_2_(50 µM), we were able to promote activation of the p53 group by overexpressing the SRXN1 enzyme, or to block the p53 group while promoting the FOXO1 group with the SRXN1 inhibitor J14.

Analysis of the RNA-seq data revealed little overlap in the genes significantly upregulated or downregulated in the p53 and FOXO1 group. Of the 881 genes significantly upregulated by either group, ∼10% were shared between the two groups. Similarly, of the ∼1200 genes downregulated by either group, there was ∼9% overlap between the two groups (**Fig. 7e**). Despite the large differences in gene expression changes between the two TF groups, Gene Set Enrichment Analysis using the Hallmark gene set revealed a large overlap in the pathways enriched in both the p53 and FOXO1 groups (**Table 1**). Specifically, of the 28 and 31 gene sets enriched in p53 and FOXO1 groups, respectively, 17 were shared and had the same sign NES. This accounts for a Jaccard overlap (intersection over union) of 40.5%. This finding supports our hypothesis that the FOXO1 partially serves as a backup mechanism to support the cell’s response to oxidative stress while translation of the p53 group is halted. However, distinct gene sets were also enriched by each TF module supporting the idea that cells also regulate separate transcriptional programs depending on the severity of H_2_O_2_ stress (**Table 2, 3**). In addition, KEGG and GO Biological Processes (GO:BP) analysis on differentially expressed genes revealed a large degree of overlap between the two groups. FOXO1 and p53 groups shared 73 GO:BP pathways and 6 KEGG pathways, corresponding to 18.6% and 23.1% overlaps, respectively (Jaccard index) (**Supp. Table 1, 4**). As in our GSEA analysis, distinct KEGG and GO:BP pathways were enriched in each TF module (**Supp. Table 2, 3 & Supp. Data 1, 2**).

**Table 1.** GSEA analysis was used to find overlapping Hallmark gene sets that were enriched in cells with either p53 active (“p53 Group”) or suppressed (“FOXO1 Group”) after H_2_O_2_ treatment. “NES” = Normalized Enrichment Score. Red color indicates a positive enrichment, and blue represents a negative enrichment.

| Gene Set | NES - p53 Group | NES - FOXO1 Group |
| --- | --- | --- |
| Apoptosis | 2.11 | 2.25 |
| DNA Repair | 1.72 | 2.06 |
| Epithelial Mesenchymal Transition | 1.59 | 1.66 |
| Estrogen Response Early | -1.64 | 1.80 |
| Estrogen Response Late | -1.67 | 1.80 |
| G2M Checkpoint | -2.68 | -1.72 |
| Glycolysis | 1.40 | 1.86 |
| Hypoxia | 2.57 | 2.97 |
| Inflammatory Response | 2.20 | 1.43 |
| Interferon Gamma Response | 2.20 | 1.76 |
| Kras Signaling Up | 1.74 | 1.43 |
| Mitotic Spindle | -2.38 | -2.56 |
| MTORC1 Signaling | 2.11 | 1.58 |
| Myc Targets V1 | -1.61 | 2.22 |
| Myc Targets V2 | -2.00 | 1.46 |
| Myogenesis | 1.52 | 1.76 |
| p53 Pathway | 3.06 | 2.76 |
| Reactive Oxygen Species Pathway | 2.07 | 2.07 |
| TNFA Signaling Via NFkB | 2.87 | 3.28 |
| UV Response Up | 1.57 | 2.29 |
| Xenobiotic Metabolism | 1.83 | 1.85 |

**Table 2.** GSEA analysis was used to find unique Hallmark gene sets that were enriched in cells with p53 active (“p53 Group”) after H_2_O_2_ treatment. “NES” = Normalized Enrichment Score. Red color indicates a positive enrichment, and blue represents a negative enrichment.

| Gene Set | NES |
| --- | --- |
| E2F Targets | -2.08 |
| Heme Metabolism | 2.14 |
| IL2 STAT5 Signaling | 1.44 |
| IL6 JAK STAT3 Signaling | 1.95 |
| Interferon Alpha Response | 1.50 |
| PI3K AKT MTOR Signaling | 1.72 |
| Unfolded Protein Response | 1.77 |

**Table 3.** GSEA analysis was used to find unique Hallmark gene sets that were enriched in cells with FOXO1 active (“FOXO1 Group”) after H_2_O_2_ treatment. “NES” = Normalized Enrichment Score. Red color indicates a positive enrichment, and blue represents a negative enrichment.

| Gene Set | NES |
| --- | --- |
| Adipogenesis | 2.00 |
| Androgen Response | -1.51 |
| Cholesterol Homeostasis | 1.63 |
| Coagulation | 1.74 |
| Complement | 1.41 |
| Fatty Acid Metabolism | 1.77 |
| Oxidative Phosphorylation | 2.89 |
| Protein Secretion | -1.76 |
| TGF Beta Signaling | -1.61 |
| UV Response DN | -1.92 |

In summary, these analyses support a model where shuttling TFs in the FOXO1 group provide a backup mechanism to transcriptionally respond to high levels of H_2_O_2_ while translation is inhibited (**Fig. 7f**). Translation attenuation therefore appears to function as a molecular switch that rewires transcription factor usage in response to increasing H_2_O_2_.

## DISCUSSION

In response to H_2_O_2_ stress, cells activate several TFs which upregulate hundreds of target genes that help to restore redox balance, repair oxidative damage, and, when damage is extreme, activate programmed cell death. We previously categorized these TFs into two groups based on their H₂O₂ dose and activation timing. The "p53 group" (p53, NRF2, ATF4, JUN) activates rapidly under low H₂O₂ stress, while the "FOXO1 group" (FOXO1, FOXO3, NF-κB, NFAT1) remains inactive. In contrast, high H₂O₂ stress elicits a biphasic response: the FOXO group activates rapidly, while the p53 group is suppressed. Eventually the FOXO1 group switches off and the p53 group is activated^9^.

Here we uncovered a switch-like response where the activity of multiple translation control pathways changed at the H_2_O_2_ dose that p53 accumulation is suppressed. Activation of the ISR through GCN2 and PERK, activation of eEF2K, and inhibition of mTORC1 signaling all coincide with loss of p53 accumulation. Strikingly, these events occur at a particular H_2_O_2_ concentration threshold (∼100 μM) where two cellular populations emerge; one with high p53 and sustained “pro-translation” signaling and another with low p53 abundance and attenuated translation. We previously observed that accumulation of NRF2, ATF4, and JUN is blocked at this same dose, suggesting that translation attenuation broadly restricts the p53 group of TFs under high levels of H_2_O_2_. Consistent with this, proteasome and ubiquitin ligase inhibitors failed to restore NRF2 and ATF4 accumulation under high H_2_O_2_ stress, despite increasing their abundance in unstressed cells. (**Supp. Fig. 1c, d & 2a, b**).

The repression of the p53 group of TFs is accompanied by rapid nuclear shuttling of FOXO1, NF-κB and NFAT1. Although the p53 and FOXO1 group of TFs regulate largely distinct target genes, both programs converge on several overlapping stress-response pathways. This switch from p53 to FOXO1 group activation at high H_2_O_2_ may therefore serve multiple functions. It could serve as a back-up mechanism, upregulating pathways critical for responding to H_2_O_2_ stress while translation is attenuated and the p53 group of TFs is repressed. Since translation is suppressed, the genes upregulated by the FOXO1 group might prime cells for recovery once translational capacity is restored. However, previous studies have revealed that while H_2_O_2_ limits translation, selective translation of specific transcripts still occurs^51,52^. Consistent with this, we found that translation (as measured by OPP) under high H_2_O_2_ is reduced but not abolished relative to cycloheximide controls suggesting that some translation is still occurring (**Fig. 3b**). Thus, the changes in gene expression might also serve to upregulate genes that can be translated despite activation of the ISR, eEF2K and suppression of mTORC1 signaling. Such preferential translation could be influenced by promoter or transcript features, as described in budding yeast during glucose starvation.

The fact that the translation inhibitory events studied here occur at approximately the same dose is striking. A major open question in the redox field is how specificity is achieved in redox signaling^53–57^. Many of the current models for signaling specificity are centered around the highly abundant family of antioxidant proteins, the peroxiredoxins (PRDXs). PRDXs contain a reactive cysteine residue 5-7 orders of magnitude more reactive towards H_2_O_2_ than other cysteines in the cell^58^. Due to their high abundance and extreme sensitivity for H_2_O_2_, PRDXs are a strong buffer against fluctuations in H_2_O_2_ levels. However, inactivation of PRDXs through hyper-oxidation of their catalytic cysteine residue or via phosphorylation can lead to a buildup of H_2_O_2_ and subsequent oxidation of proteins in the vicinity^59–61^. Several studies have also shown that once oxidized, PRDXs can transfer their oxidation state to downstream binding partners altering their activity^62–64^. An attractive possibility is that PRDX-mediated redox sensing converges on multiple translation control hubs, thereby coordinating translational repression with TF switching at a defined H_2_O_2_ threshold. Indeed, we have previously shown that genetic and chemical perturbations of PRDXI/II alter the doses at which certain TFs are activated in response to H_2_O_2_ further supporting this model^9^. However, although these models seem to be true for some cysteine oxidation events, there is recent evidence for alternative modes of protein oxidation which rely on oxidation of L-tryptophan, the presence of bicarbonate, covalent addition of Coenzyme A, or the covalent addition of a peroxiredoxin protein termed “peroxiredoxinylation”^65–68^. Given this, one limitation of our study is that we did not reveal how the oxidative signal reaches each translation control hub at approximately the same dose of H_2_O_2_. We believe this leaves an interesting avenue to explore.

## ACKNOWLEDGEMENTS

We thank members of the Paek lab and the Thorne lab for helpful comments and discussion. We thank members of C.P. graduate committee (A. Capaldi, R. Buchan, M. Padi, and T. Weinert) for advice on experiments and discussion of data. This work was supported by National Institutes of Health Grants RO1-GM130864 (A.L.P.), R35GM158008 (A.L.P.), NIH T32-GM136536 (C.P.), NIH T32-GM132008 (W.M.-S.).

## RESOURCE AVAILABILITY

### Lead contact

Requests for further information and resources should be directed to and will be fulfilled by the lead contact, Andrew L. Paek.

### Materials availability

All unique/stable reagents generated in this study are available from the lead contact without restriction.

### Data and code availability

All custom written scripts described in the manuscript are available upon request.

## AUTHOR CONTRIBUTIONS

C.P. and A.L.P. designed the research. C.P. performed and/or contributed to all experiments. W.M-S. analyzed all the data from Bulk RNA sequencing. E.J. contributed to experiments from figure 1. C.P. and L.S. contributed to making the fluorescent tagged lines and knockout lines. C.P., W.M.S., and A.L.P. performed data analysis. C.P. and A.L.P. wrote the manuscript. A.L.P., C.P., W.M-S. acquired funding. All authors have reviewed and commented on the manuscript.

## METHODS

### Cell lines

MCF7 cells were a gift from Galit Lahav, Harvard Medical School and were validated by short tandem repeat profiling by the University of Arizona Genetics Core in 2019. U-2 OS (HTB-96) and MCF10A (CRL-10317) cells were obtained from ATCC and were validated by ATCC using short tandem, repeat profiling in 2018. All cell lines were tested free of mycoplasma by DAPI stain.

### Cell culture

MCF7 cells were grown in Roswell Park Memorial Institute 1640 medium (RPMI) supplemented with 10% FBS, 100 units/mL penicillin, 100 μg/mL streptomycin, and 25 ng/mL amphotericin B. U-2 OS (HTB-96) were grown in Dulbecco’s Modified Eagle Medium (DMEM) supplemented with the same concentrations of FBS and antibiotics as mentioned above. MCF10A (CRL-10317) were grown in DMEM/F-12 (Invitrogen #11330-032) media supplemented with 5% Horse serum (Invitrogen#16050-122), EGF (20 ng/mL final), Hydrocortisone (0.5 mg/mL final), Cholera Toxin (100 ng/mL final), Insulin (10 μg/mL final), 1% Pen/Strep (100x solution, Invitrogen #15070-063).

### Plasmid and cell line construction

Lentivirus was produced and infection was carried out as described by ref. (Tiscornia, G., et al., 2006)^57,69^. On day 1, 5*10^6^ HEK 293T cells were plated into 10cm dishes in DMEM + 10% FBS. After 24h, media was replaced with 12mL of fresh DMEM + 10% FBS + 10mM HEPES. The transfection reagent was prepared by adding 18µl TransIT Transfection Reagent LT1 (Cat# MIR 2304) to 800µl of serum-free DMEM. 3.2µg of the lentiviral plasmid, 1.8µg of pMDLg/pRRE (Addgene: 12251), 0.7µg of pRSV-Rev (Addgene: 12253) and 0.3µg of pMD2.G (Addgene: 12259) were then mixed and incubated at room temperature for 30min and then added to the 293T cells. On Day 4, the media was harvested and stored in a 50mL falcon tube and stored overnight at 4°C. 5 mL of fresh DMEM + 10% FBS + 10mM HEPES was added. On day 5, the remaining media was harvested to the falcon tube. This was then spun down at 800g for 20 min and filtered through a 0.45 µM filter. Virus was stored at −80°C in 1 ml aliquots.

For lentiviral infection we plated 5*10^4^ cells on day one. After 24h, we prepared lentivirus by adding protamine sulfate to a final concentration of 8µg/mL and 1µl of 1M HEPES to 1mL of thawed virus. Media was then aspirated from cells and replaced with 1mL of virus and another 1mL of fresh cell culture media. Cells were incubated at 37°C for 4–6 h and then viral media was replaced with 10mL of fresh tissue culture media.

H2B-ECFP (pRRLH-UbCpp-H2B-CFP)^9,70^ and p53-mScarleti3 (pLV-Puro-hPGK-p53-mScarlet-I3; VectorBuilder ID: VB250627-1353ana) lentiviral vectors were generated as described above. pLenti-FoxO1-Clover was a gift from Peter Rotwein (Addgene plasmid # 67759; http://n2t.net/addgene:67759; RRID: Addgene_67759)^71^. MCF7 cells were infected as described above after vector preparation and selected with hygromycin and/or puromycin.

To generate knockout MCF7 cell lines, Mammalian CRISPR Lentiviral Vector (Single gRNA) vectors were designed and ordered from https://en.vectorbuilder.com/. For the eEF2K (*EEF2K*) knockout, the vector ID is VB900141-4525xws. For the 4EBP1 (*EIF4EBP1*) knockout, the vector ID is VB250121-1173xdy. For TSC2 (*TSC2*), the vector ID is VB900145-4201ddj. Vectors were used to generate lentivirus as described above. Cells were selected with puromycin, and knockouts were validated with western blot.

pMRX-IP-GFP-LC3-RFP-LC3ΔG was a gift from Noboru Mizushima (Addgene plasmid # 84572; http://n2t.net/addgene:84572; RRID: Addgene_84572). This plasmid was used to generate a retrovirus containing the autophagic flux reporter as seen in (Supp. Fig. 1d, e). Retrovirus production and infection follow the same protocol as lentivirus above except for these changes to the packaging plasmids: 3.2µg of the retroviral plasmid, 1.8µg of pUMVC (Addgene: 8449) and 0.3µg of pCMV-VSV-G (Addgene: 8454) were then mixed and incubated at room temperature for 30min and then added to the 293T cells. Infected cells were selected with puromycin.

### Cell treatments

For H_2_O_2_ treatments, H_2_O_2_ was diluted in PBS then added directly to the newly replaced media to get the final concentrations indicated in each experiment. All IF experiments were treated with H_2_O_2_ for 3hrs before fixation. The stock of H_2_O_2_ (Sigma Aldrich H1009-100ml) was replaced every 2 months. Cells were pretreated with Nutlin-3a (Fig. 1a), MG132 (Fig. 1b, Supp. Fig. 1c, d), Chloroquine (CQ) and Bafilomycin A1 (BafA1) (Fig. 2c), GCN2iB (GCN2i) and GSK2656157 (PERKi) (Fig. 5, Supp. Fig. 4a-c), KU60019 (ATMi) (Supp. Fig. 1a, b), Actinomycin D (ActD) (Supp. Fig. 1i), and Cycloheximide (CHX) (Supp. Fig. 1j) for 10 minutes prior to H_2_O_2_ treatment. Cells were pretreated with Nelfinavir (NFR) (Supp. Fig. 5a), NH125 (Supp. Fig. 5b) Rapamycin (Fig. 6a, Supp. Fig. 5d, e), and AZD8055 (Fig. 6b, Supp. Fig. 5d, e) for 2 hours prior to H_2_O_2_ treatment. Cells were pretreated with sodium arsenite (NaAsO_2_) (Supp. Fig. 4c, d) for 30 minutes prior to H_2_O_2_ treatment or fixation, respectively.

### Quantitative PCR

Total RNA was isolated from cell lysates using the Qiagen RNeasy Extraction Kit (Cat: 23227) according to the manufacturer’s instructions. The isolated RNA was reverse transcribed into cDNA using the Applied Biosystems High-Capacity Reverse Transcription Kit (Cat: 43-688-14). Quantitative real-time PCR was performed using Applied Biosystems Real-Time PCR System with Applied Biosystems SYBR Green Universal Master Mix (Cat:43-091-55). The relative expression levels of target genes were normalized to the expression level of GAPDH, which was used as an internal control, and calculated the 2-ΔΔCT method^72^. Primer sequences are listed in Methods Table 1.

### O-propargyl-puromycin incorporation assay

The Click-&-Go® Plus 647 OPP Protein Synthesis Assay Kit (Vector Laboratories #CCT-1496) was used as instructed by the manufacturer. Briefly, 1hr before cell fixation CHX was added to appropriate wells. Then, 30min until cell fixation OPP (20µM) was added to all wells. After 30 minutes, cells were washed once with PBS then fixed (PBS +3.7% formaldehyde) and then permeabilized (PBS +0.1% Triton X-100). A click chemistry reaction was then performed to attach the far-red fluorescent AZDye 647 Azide Plus probe to OPP. OPP intensity was measured via fluorescent microscopy.

### Western blots

Cells (approx.100,000 cells/dish) were plated to a 6cm dish and incubated for 48h. They were washed with PBS, scraped off the plate, centrifuged (5min, 3K rpm, room temperature) and the cell pellet was lysed using a Lysis Buffer (25 mM Tris pH 7.6, 150 mM NaCl, 1% NP-40, 1% Na-deoxycholate, and 0.1% SDS in water + protease inhibitor cocktail [Sigma-Aldrich: P8340] + phosphatase inhibitor cocktail [Sigma-Aldrich: P0044] + okadaic acid [Thermo Scientific: J60155.I&) + sodium fluoride). The cells were spun down (30min, 14K rpm, 4°C) and the supernatant was used to measure protein concentration by using a BCA assay [Thermo Scientific: 23227]. Equal protein concentrations were loaded onto a NuPAGE 4-12% Bis-Tris gels [Invitrogen: NP0322BOX]. Protein was transferred to a PVDF membrane using the iBlot™ 2 system – Invitrogen: IB21001, IB24002. The PVDF membrane was then incubated in blocking solution (PBS, 5% BSA, 0.1% Tween 20) for 1 h at room temperature. The membrane was incubated with primary antibodies at 4°C overnight, rinsed three times with PBST and incubated with secondary antibodies for 1 h at room temperature. The blot was imaged on the LI-COR Odyssey.

Primary Antibodies used: Anti-β-Actin (ACTB) mouse mAb Cat#A2228 clone AC-74 (Sigma) was used as a loading control (1:20,000). Anti-eEF2K rabbit pAb from Cell Signaling Cat# 3692S (1:1000). Anti-Tuberin/TSC2 (D93F12) XP® rabbit mAb from Cell Signaling Cat# 4308S (1:1000). Anti-4E-BP1 (53H11) rabbit mAb from Cell Signaling Cat# 9644S (1:1000).

Secondary antibodies used: IRDye® 800CW Goat anti-Mouse IgG from LICORbio Cat# 926-32210 (1:20,000). Mouse anti-rabbit IgG-HRP from Santa Cruz Cat# sc-2357 (1:25,000). For visualizing the HRP signal, the SuperSignal™ West Pico PLUS Chemiluminescent Substrate kit from Thermo Scientific Cat# 34580 was used as instructed by the manufacturer.

### Live cell microscopy

Autophagy Movie (Fig. 2) – Cells (20,000 cells/well) were plated on 24 well glass bottom plates (CellVis Cat# P24-1.5H-N) which were coated with poly L-lysine (Sigma Cat# P4707) and allowed to grow for 72hrs. Cells were grown in the appropriate media as mentioned above and then rinsed with PBS and given DMEM Fluorobrite media (Gibco Cat #A18967-01) with 2% FBS,100 units/mL penicillin, 100 μg/mL streptomycin, 25 ng/mL amphotericin B, and 1x GlutaMAX (Thermo Scientific Cat# 35050061). Cells were imaged every 30min for 24hrs by a Nikon Eclipse Ti-E microscope. Data was acquired using the NIS Elements AR 5.21.02 software. A thin layer of mineral oil was used to prevent media evaporation. Temperature (37 °C) and 5% CO2 levels were maintained using the OKO labs incubation system. H2B-iRFP was imaged using the (Chroma) ET-Cy5.5 filter set (exposure time = 400ms). GFP-LC3 was imaged using the AT-EGFP/F Filter Set (exposure time = 400ms). RFP-LC3ΔG was imaged using the AT-TRITC Filter Set (exposure time = 400ms). Bafilomycin A1 or Chloroquine were added to cells 10 minutes prior to H_2_O_2_ or rapamycin treatment.

P53 Movie (Fig. 3) – Cells (35,000 cells/well) were plated on 24 well glass bottom plates (CellVis Cat# P24-1.5H-N) which were coated with poly L-lysine (Sigma Cat# P4707) and allowed to grow for 48hrs. Cells were grown in the appropriate media as mentioned above and then rinsed with PBS and given DMEM Fluorobrite media (Gibco Cat #A18967-01) with 2% FBS,100 units/mL penicillin, 100 μg/mL streptomycin, 25 ng/mL amphotericin B, and 1x GlutaMAX (Thermo Scientific Cat# 35050061). Cells were imaged every 20min for 8hrs by a Nikon Eclipse Ti-E microscope. Data was acquired using the NIS Elements AR 5.21.02 software. A thin layer of mineral oil was used to prevent media evaporation. Temperature (37 °C) and 5% CO2 levels were maintained using the OKO labs incubation system. H2B-CFP was imaged using the C-FL AT ECFP/Cerulean Filter Set (Chroma) (exposure time = 400ms). P53-mScarlet-I3 was imaged using the AT-TR/mCH Filter Set (Chroma) (exposure time = 200ms).

### Live cell analysis

Single cell tracking and segmentation - Cells were automatically tracked using the code at https://github.com/oylab/oyLabImaging. Segmentation was performed using StarDist, with LAP tracking for cell traces. Traces were filtered by cell area (cells below a threshold of 200 pixels were discarded).

Automatic death labeling - Morphological features per cell per frame were fed through a feed forward neural network to predict a frame-wise death/life state. Full track predictions are then passed through a Hidden Markov Model for correction and final death prediction. Predictions are evaluated for accuracy visually and corrected as necessary. (Method/software subject of manuscript in prep).

Autophagy analysis - GFP/RFP ratios (LC3/LC3ΔG) were calculated for each cell, as total GFP/total RFP. Lower ratio means more autophagy. This ratio was averaged across all cells per treatment and visualized by normalizing the full trace to the first time point values, with standard error values derived from the experimental distribution at each frame. The derivative graph was calculated by differencing adjacent time points for each cell, then averaging the traces for each experiment. These results were then smoothed using a centered rolling window of three frames.

p53 movie analysis (Fig. 3) – Using a custom MATLAB script, mean fluorescent intensity of p53-mScarlet-I3 was measured at each timepoint, with standard error values derived from the experimental distribution at each frame. Briefly, fluorescent channels were background subtracted using the tophat transformation, nuclei were then segmented using the H2B-CFP channel and the CellPose “nuclei” model with an average cell diameter of 20, and finally regionprops was used to extract mean intensities per fluorescent channel. A custom MATLAB script was then used to plot mean p53-mScarlet-i3 traces normalized to the starting value. The smoothed derivative graph was calculated by first applying a Savitzky-Golay filter (window size = 5 frames, polynomial order = 2) to the log2-normalized traces. Then the derivative was calculated by differencing adjacent time points. Finally, a moving average filter of 5 frames was used to further smooth the resulting derivative traces.

### Immunofluorescence

Cells (3500 cells/well) were plated on 96-well plates from Greiner Bio-One (MicroClear 96 well Cat #655090-99). Cells were allowed to grow for 48hrs prior to treatment. Cells were fixed with 2% PFA (Thermofisher Cat #043368.9M) for 10 min then washed twice in PBS. Cells were then permeabilized using 0.1% Triton X-100 in PBS for 10min then washed with PBS twice. Cells were then blocked with 2% BSA in PBS for 30 min and then incubated overnight at 4°C with primary antibodies diluted in 2% BSA and 0.1% Tween in PBS. Cells were then washed two times with PBS followed by incubation with secondary antibodies in 2% BSA and 0.1% Tween at room temperature for 1 hour. The cells were then washed two times in PBS, counterstained with DAPI, then imaged in PBS. Images were analyzed using Cell Profiler^73^. To extract nuclear intensities, nuclei were segmented using DAPI. To obtain cytoplasmic intensities, a ring of 3 pixels wide was drawn around the nuclear mask and mean cytoplasmic levels were extracted using this mask. To obtain cellular intensities, the nuclear and cytoplasmic regions were added together, and mean intensities were calculated using this region. All plots were made using MATLAB version R2025a.

Primary antibodies used: Anti-p53 (DO-1) mouse mAb from Santa Cruz Cat# sc-126 (1:500). Anti-Phospho-Chk2 (Thr68) rabbit pAb from Cell Signaling Cat# 2661S (1:200). Anti-Phospho-eIF2a (S51) (119A11) rabbit mAb from Cell Signaling Cat# 3597S (1:200). Anti-Phospho-S6 Ribosomal Protein (S235/236) rabbit mAb from Cell Signaling Cat# 2211S (1:500). Anti-Phospho-4E-BP1 (Thr37/46) rabbit mAb from Cell Signaling Cat# 2855S (1:500). Anti-Phospho-eEF2 (Thr56) rabbit pAb from Cell Signaling Cat# 2331S (1:500). Anti-ATF4 (D4B8) rabbit mAb from Cell Signaling Cat# 11815S (1:200). Anti-NRF2 (D1Z9C) XP® rabbit mAb from Cell Signaling Cat# 12721S (1:500). Anti-LC3B (D11) XP® rabbit mAb from Cell Signaling Cat# 3868S (1:500). Anti-G3BP mAb from BD Biosciences Cat# 611126 (1:1000).

Secondary antibodies used: Goat Anti-Rabbit IgG (H+L) Alexa Fluor™ 488 Cell Signaling Cat# A-11034 (1:500). Goat anti-Mouse IgG (H+L) Alexa Fluor™ 594 Cat# A-11032 (1:500). Goat anti-Mouse IgG (H+L) Alexa Fluor™ 488 (1:500) Cat# A-11001. Goat anti-Rabbit IgG (H+L) Highly Cross-Adsorbed Secondary Antibody, Alexa Fluor™ 568 (1:500) Cat# A-11036.

Cells were imaged on a Nikon Eclipse Ti-E microscope. Data was acquired using the NIS Elements AR 5.21.02 software and visualized with NIS Elements Viewer 5.21. DAPI was imaged using the AT-DAPI Filter Set (exposure time = 10-20ms). Alexa Fluor 488 was imaged using AT-EGFP/F Filter Set (exposure time = 50-200ms). Alexa Fluor 594 and mCherry were imaged using the AT mCherry Filter Set (exposure time = 20-200ms). AZDye647 was imaged using the (Chroma) ET-Cy5.5 filter set (exposure time = 20ms). Alexa Fluor 568 was imaged using FOXO1-Clover was imaged using the AT mCherry Filter Set (exposure time = 50-200ms). FOXO1-Clover was imaged using AT-EGFP/F Filter Set (exposure time = 100ms).

FOXO1-Clover images were background subtracted in MATLAB using the “imbackground” function from the p53CinemaManual package^74^ which applies morphological operations using two structuring elements. Default structuring element sizes were 10 and 100 pixels which are optimized for images acquired using a 20x objective. Cells not expressing the FOXO1-Clover reporter were thrown out of the analysis in Figure 7 based on cellular intensity thresholding using the MATLAB function “multithresh” (Otsu’s method) to generate 2 threshold values.

### Differential expression analysis on Bulk RNA-seq data

Bulk RNA sequencing data was taken from a study on the temporal dynamics of transcription factor response to H_2_O_2_ stress^9^. Cells overexpressing Sulfiredoxin or treated with J14 (a Sulfiredoxin inhibitor) were treated with 50 µM H_2_O_2_ leading to p53 or FOXO1 activation, respectively. Counts from bulk sequencing data were extracted using the Rsubread package^75^ with hg38 annotation. The DESeq2 package in R^76^ was used to compare cells overexpressing Sulfiredoxin treated with H_2_O_2_ (0 µM vs. 50 µM). The Wald statistic from these results was used for gene set enrichment analysis (GSEA) using the Hallmark gene set and the clusterProfiler R package^77^. The same analysis was performed for cells treated with J14 and H_2_O_2_ (0 µM vs. 50 µM). KEGG and GO analysis were performed using clusterProfiler for each DESeq2 result dataset, with genes determined to be significant with an absolute log2 fold change greater than 1 and an adjusted p-value less than 0.01. For each gene set analysis, shared hits with an adjusted p-value less than 0.05 were found between the J14 treated and Sulfiredoxin overexpression group. Overlap between enriched GSEA, KEGG, and GO pathway sets was quantified using the Jaccard index, defined as the size of the intersection divided by the size of the union of pathways between groups.

For the Venn diagram in Supplemental Figure 6c, upregulated genes from both the J14 (i.e. – FOXO1 group) and Sulfiredoxin overexpression (i.e. – p53 group) groups are defined as those having a log2 fold change greater than 0.2 and adjusted p-value less than 1e-5. Downregulated genes from both the J14 (i.e. – FOXO1 group) and Sulfiredoxin overexpression (i.e. – p53 group) groups are defined as those having a log2 fold change less than 0.2 and adjusted p-value less than 1e-5. Unique and similar genes were found from these lists using the functions “setdiff” and “intersect” in MATLAB, respectively.

### Statistics and reproducibility

No statistical methods were employed to predetermine sample size. The experiments were not randomized, and the investigators were not blinded to the source of the experimental samples and outcome assessment.

All immunofluorescence experiments were replicated a minimum of three times with reproducible results. P-values were acquired by first performing a two-sample t-test then adjusted for multiple hypotheses using the Benjamini-Hochberg procedure (* ≤ 0.05, ** ≤ 0.01, *** ≤ 0.001). Spearman’s rank correlation was used to assess the monotonic relationship between log2 nuclear p53 and log2 cellular OPP intensity in Supp. Fig. 1k. Time lapse movies were run at least three times with reproducible results. Cell number was maintained as the impact of H_2_O_2_ exposure decreases with cell number^9^.

**Methods Table 1:**
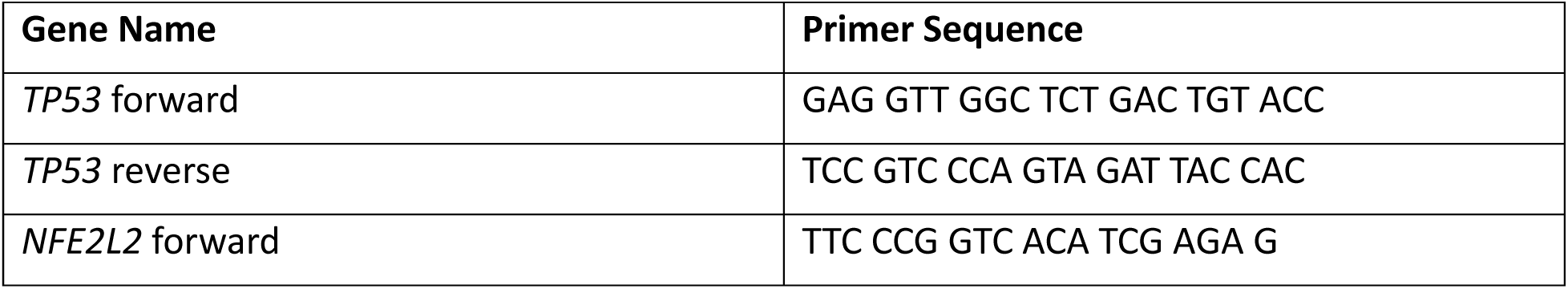
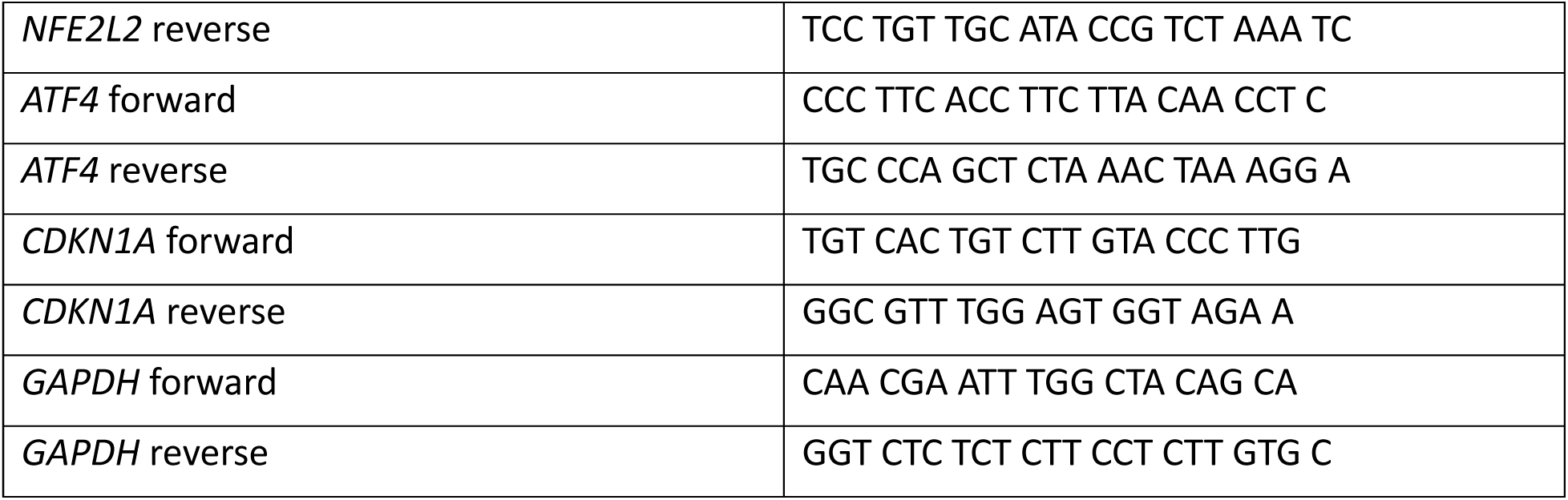
Forward and Reverse primers used for qPCR.

## REFERENCES

1. Niethammer, P., Grabher, C., Look, A.T., and Mitchison, T.J. (2009). A tissue-scale gradient of hydrogen peroxide mediates rapid wound detection in zebrafish. Nature 459, 996–999. 10.1038/nature08119.

2. Sundaresan, M., Yu, Z.-X., Ferrans, V.J., Irani, K., and Finkel, T. (1995). Requirement for Generation of H2O2 for Platelet-Derived Growth Factor Signal Transduction. Science 270, 296–299. 10.1126/science.270.5234.296.

3. Rampon, C., Volovitch, M., Joliot, A., and Vriz, S. (2018). Hydrogen Peroxide and Redox Regulation of Developments. Antioxidants (Basel) 7, 159. 10.3390/antiox7110159.

4. Keyer, K., and Imlay, J.A. (1996). Superoxide accelerates DNA damage by elevating free-iron levels. Proceedings of the National Academy of Sciences 93, 13635–13640. 10.1073/pnas.93.24.13635.

5. Shigenaga, M.K., Hagen, T.M., and Ames, B.N. (1994). Oxidative damage and mitochondrial decay in aging. Proc Natl Acad Sci U S A 91, 10771–10778. 10.1073/pnas.91.23.10771.

6. Mylonas, C., and Kouretas, D. (1999). Lipid peroxidation and tissue damage. In Vivo 13, 295–309.

7. Lisanti, M.P., Martinez-Outschoorn, U.E., Lin, Z., Pavlides, S., Whitaker-Menezes, D., Pestell, R.G., Howell, A., and Sotgia, F. (2011). Hydrogen peroxide fuels aging, inflammation, cancer metabolism and metastasis. Cell Cycle 10, 2440–2449. 10.4161/cc.10.15.16870.

8. Marinho, H.S., Real, C., Cyrne, L., Soares, H., and Antunes, F. (2014). Hydrogen peroxide sensing, signaling and regulation of transcription factors. Redox Biology 2, 535–562. 10.1016/j.redox.2014.02.006.

9. Jose, E., March-Steinman, W., Wilson, B.A., Shanks, L., Parkinson, C., Alvarado-Cruz, I., Sweasy, J.B., and Paek, A.L. (2024). Temporal coordination of the transcription factor response to H2O2 stress. Nat Commun 15, 3440. 10.1038/s41467-024-47837-w.

10. Hafner, A., Bulyk, M.L., Jambhekar, A., and Lahav, G. (2019). The multiple mechanisms that regulate p53 activity and cell fate. Nat Rev Mol Cell Biol 20, 199–210. 10.1038/s41580-019-0110-x.

11. Nadeau, P.J., Charette, S.J., Toledano, M.B., and Landry, J. (2007). Disulfide Bond-mediated Multimerization of Ask1 and Its Reduction by Thioredoxin-1 Regulate H2O2-induced c-Jun NH2-terminal Kinase Activation and Apoptosis. MBoC 18, 3903–3913. 10.1091/mbc.e07-05-0491.

12. Shi, T., van Soest, D.M.K., Polderman, P.E., Burgering, B.M.T., and Dansen, T.B. (2021). DNA damage and oxidant stress activate p53 through differential upstream signaling pathways. Free Radical Biology and Medicine 172, 298–311. 10.1016/j.freeradbiomed.2021.06.013.

13. Hanson, R.L., and Batchelor, E. (2022). Coordination of MAPK and p53 dynamics in the cellular responses to DNA damage and oxidative stress. Molecular Systems Biology 18, e11401. 10.15252/msb.202211401.

14. Bulavin, D.V., Saito, S., Hollander, M.C., Sakaguchi, K., Anderson, C.W., Appella, E., and Fornace, A.J. (1999). Phosphorylation of human p53 by p38 kinase coordinates N-terminal phosphorylation and apoptosis in response to UV radiation. The EMBO Journal 18, 6845–6854. 10.1093/emboj/18.23.6845.

15. Milne, D.M., Campbell, L.E., Campbell, D.G., and Meek, D.W. (1995). p53 Is Phosphorylated in Vitro and in Vivo by an Ultraviolet Radiation-induced Protein Kinase Characteristic of the c-Jun Kinase, JNK1 (∗). Journal of Biological Chemistry 270, 5511–5518. 10.1074/jbc.270.10.5511.

16. Uziel, T., Lerenthal, Y., Moyal, L., Andegeko, Y., Mittelman, L., and Shiloh, Y. (2003). Requirement of the MRN complex for ATM activation by DNA damage. EMBO J 22, 5612–5621. 10.1093/emboj/cdg541.

17. Stommel, J.M., and Wahl, G.M. (2004). Accelerated MDM2 auto-degradation induced by DNA-damage kinases is required for p53 activation. The EMBO Journal 23, 1547–1556. 10.1038/sj.emboj.7600145.

18. Guo, Z., Kozlov, S., Lavin, M.F., Person, M.D., and Paull, T.T. (2010). ATM Activation by Oxidative Stress. Science 330, 517–521. 10.1126/science.1192912.

19. Buttgereit, F., and Brand, M.D. (1995). A hierarchy of ATP-consuming processes in mammalian cells. Biochemical Journal 312, 163–167. 10.1042/bj3120163.

20. Maechler, P., Jornot, L., and Wollheim, C.B. (1999). Hydrogen Peroxide Alters Mitochondrial Activation and Insulin Secretion in Pancreatic Beta Cells*. Journal of Biological Chemistry 274, 27905–27913. 10.1074/jbc.274.39.27905.

21. LaCagnin, L.B., Bowman, L., Ma, J.Y., and Miles, P.R. (1990). Metabolic changes in alveolar type II cells after exposure to hydrogen peroxide. American Journal of Physiology-Lung Cellular and Molecular Physiology 259, L57–L65. 10.1152/ajplung.1990.259.2.L57.

22. Gebauer, F., and Hentze, M.W. (2004). Molecular mechanisms of translational control. Nat Rev Mol Cell Biol 5, 827–835. 10.1038/nrm1488.

23. Ryazanov, A.G., Shestakova, E.A., and Natapov, P.G. (1988). Phosphorylation of elongation factor 2 by EF-2 kinase affects rate of translation. Nature 334, 170–173. 10.1038/334170a0.

24. Shenton, D., Smirnova, J.B., Selley, J.N., Carroll, K., Hubbard, S.J., Pavitt, G.D., Ashe, M.P., and Grant, C.M. (2006). Global translational responses to oxidative stress impact upon multiple levels of protein synthesis. J Biol Chem 281, 29011–29021. 10.1074/jbc.M601545200.

25. Sanchez, M., Lin, Y., Yang, C.-C., McQuary, P., Rosa Campos, A., Aza Blanc, P., and Wolf, D.A. (2019). Cross Talk between eIF2α and eEF2 Phosphorylation Pathways Optimizes Translational Arrest in Response to Oxidative Stress. iScience 20, 466–480. 10.1016/j.isci.2019.09.031.

26. Ma, X.M., and Blenis, J. (2009). Molecular mechanisms of mTOR-mediated translational control. Nat Rev Mol Cell Biol 10, 307–318. 10.1038/nrm2672.

27. Chen, L., Xu, B., Liu, L., Luo, Y., Yin, J., Zhou, H., Chen, W., Shen, T., Han, X., and Huang, S. (2010). Hydrogen peroxide inhibits mTOR signaling by activation of AMPKalpha leading to apoptosis of neuronal cells. Lab Invest 90, 762–773. 10.1038/labinvest.2010.36.

28. Erales, J., and Coffino, P. (2014). Ubiquitin-independent proteasomal degradation. Biochim Biophys Acta 1843, 216–221. 10.1016/j.bbamcr.2013.05.008.

29. Lassot, I., Ségéral, E., Berlioz-Torrent, C., Durand, H., Groussin, L., Hai, T., Benarous, R., and Margottin-Goguet, F. (2001). ATF4 degradation relies on a phosphorylation-dependent interaction with the SCF(betaTrCP) ubiquitin ligase. Mol Cell Biol 21, 2192–2202. 10.1128/MCB.21.6.2192-2202.2001.

30. Mauthe, M., Orhon, I., Rocchi, C., Zhou, X., Luhr, M., Hijlkema, K.-J., Coppes, R.P., Engedal, N., Mari, M., and Reggiori, F. (2018). Chloroquine inhibits autophagic flux by decreasing autophagosome-lysosome fusion. Autophagy 14, 1435–1455. 10.1080/15548627.2018.1474314.

31. Kaizuka, T., Morishita, H., Hama, Y., Tsukamoto, S., Matsui, T., Toyota, Y., Kodama, A., Ishihara, T., Mizushima, T., and Mizushima, N. (2016). An Autophagic Flux Probe that Releases an Internal Control. Molecular Cell 64, 835–849. 10.1016/j.molcel.2016.09.037.

32. Lee, S.C., Zhang, J., Strom, J., Yang, D., Dinh, T.N., Kappeler, K., and Chen, Q.M. (2017). G-Quadruplex in the NRF2 mRNA 5’ Untranslated Region Regulates De Novo NRF2 Protein Translation under Oxidative Stress. Mol Cell Biol 37, e00122–16. 10.1128/MCB.00122-16.

33. Liu, J., Xu, Y., Stoleru, D., and Salic, A. (2012). Imaging protein synthesis in cells and tissues with an alkyne analog of puromycin. Proceedings of the National Academy of Sciences 109, 413–418. 10.1073/pnas.1111561108.

34. mScarlet3: a brilliant and fast-maturing red fluorescent protein | Nature Methods https://www.nature.com/articles/s41592-023-01809-y.

35. Taniuchi, S., Miyake, M., Tsugawa, K., Oyadomari, M., and Oyadomari, S. (2016). Integrated stress response of vertebrates is regulated by four eIF2α kinases. Sci Rep 6, 32886. 10.1038/srep32886.

36. Błaszczyk, L., and Ciesiołka, J. (2011). Secondary structure and the role in translation initiation of the 5’-terminal region of p53 mRNA. Biochemistry 50, 7080–7092. 10.1021/bi200659b.

37. Ferguson, A., Wang, L., Altman, R.B., Terry, D.S., Juette, M.F., Burnett, B.J., Alejo, J.L., Dass, R.A., Parks, M.M., Vincent, C.T., et al. (2015). Functional Dynamics within the Human Ribosome Regulate the Rate of Active Protein Synthesis. Molecular Cell 60, 475–486. 10.1016/j.molcel.2015.09.013.

38. Ryazanov, A.G. (1987). Ca2+/calmodulin-dependent phosphorylation of elongation factor 2. FEBS Letters 214, 331–334. 10.1016/0014-5793(87)80081-9.

39. Boyce, M., Bryant, K.F., Jousse, C., Long, K., Harding, H.P., Scheuner, D., Kaufman, R.J., Ma, D., Coen, D.M., Ron, D., et al. (2005). A Selective Inhibitor of eIF2α Dephosphorylation Protects Cells from ER Stress. Science 307, 935–939. 10.1126/science.1101902.

40. Inoki, K., Li, Y., Zhu, T., Wu, J., and Guan, K.-L. (2002). TSC2 is phosphorylated and inhibited by Akt and suppresses mTOR signalling. Nat Cell Biol 4, 648–657. 10.1038/ncb839.

41. Zhang, H., Cicchetti, G., Onda, H., Koon, H.B., Asrican, K., Bajraszewski, N., Vazquez, F., Carpenter, C.L., and Kwiatkowski, D.J. (2003). Loss of Tsc1/Tsc2 activates mTOR and disrupts PI3K-Akt signaling through downregulation of PDGFR. J Clin Invest 112, 1223–1233. 10.1172/JCI17222.

42. Qin, X., Jiang, B., and Zhang, Y. (2016). 4E-BP1, a multifactor regulated multifunctional protein. Cell Cycle 15, 781–786. 10.1080/15384101.2016.1151581.

43. Thoreen, C.C., Kang, S.A., Chang, J.W., Liu, Q., Zhang, J., Gao, Y., Reichling, L.J., Sim, T., Sabatini, D.M., and Gray, N.S. (2009). An ATP-competitive mammalian target of rapamycin inhibitor reveals rapamycin-resistant functions of mTORC1. J Biol Chem 284, 8023–8032. 10.1074/jbc.M900301200.

44. De Gassart, A., Demaria, O., Panes, R., Zaffalon, L., Ryazanov, A.G., Gilliet, M., and Martinon, F. (2016). Pharmacological eEF2K activation promotes cell death and inhibits cancer progression. EMBO reports 17, 1471–1484. 10.15252/embr.201642194.

45. Chen, Z., Gopalakrishnan, S.M., Bui, M.-H., Soni, N.B., Warrior, U., Johnson, E.F., Donnelly, J.B., and Glaser, K.B. (2011). 1-Benzyl-3-cetyl-2-methylimidazolium iodide (NH125) Induces Phosphorylation of Eukaryotic Elongation Factor-2 (eEF2): A CAUTIONARY NOTE ON THE ANTICANCER MECHANISM OF AN eEF2 KINASE INHIBITOR. Journal of Biological Chemistry 286, 43951–43958. 10.1074/jbc.M111.301291.

46. Hafner, A., Bulyk, M.L., Jambhekar, A., and Lahav, G. (2019). The multiple mechanisms that regulate p53 activity and cell fate. Nat Rev Mol Cell Biol 20, 199–210. 10.1038/s41580-019-0110-x.

47. Suzuki, T., Muramatsu, A., Saito, R., Iso, T., Shibata, T., Kuwata, K., Kawaguchi, S., Iwawaki, T., Adachi, S., Suda, H., et al. (2019). Molecular Mechanism of Cellular Oxidative Stress Sensing by Keap1. Cell Reports 28, 746–758.e4. 10.1016/j.celrep.2019.06.047.

48. Purdom-Dickinson, S.E., Sheveleva, E.V., Sun, H., and Chen, Q.M. (2007). Translational Control of Nrf2 Protein in Activation of Antioxidant Response by Oxidants. Molecular Pharmacology 72, 1074–1081. 10.1124/mol.107.035360.

49. Dey, S., Baird, T.D., Zhou, D., Palam, L.R., Spandau, D.F., and Wek, R.C. (2010). Both Transcriptional Regulation and Translational Control of ATF4 Are Central to the Integrated Stress Response. J Biol Chem 285, 33165–33174. 10.1074/jbc.M110.167213.

50. Vesely, P.W., Staber, P.B., Hoefler, G., and Kenner, L. (2009). Translational regulation mechanisms of AP-1 proteins. Mutation Research/Reviews in Mutation Research 682, 7–12. 10.1016/j.mrrev.2009.01.001.

51. Gerashchenko, M.V., Lobanov, A.V., and Gladyshev, V.N. (2012). Genome-wide ribosome profiling reveals complex translational regulation in response to oxidative stress. Proc Natl Acad Sci U S A 109, 17394–17399. 10.1073/pnas.1120799109.

52. Bruno, I., Perrucci, C., Tomè, G., Susin, G., Mazzalai, S., Sevegnani, M., Piano, A.D., Donini, L., Basso, M., Clamer, M., et al. (2025). Empowering multiplexed ultra-throughout ribosome profiling with RiboWich. Preprint at bioRxiv, 10.1101/2025.10.17.683053 https://doi.org/10.1101/2025.10.17.683053.

53. Winterbourn, C.C., and Hampton, M.B. (2008). Thiol chemistry and specificity in redox signaling. Free Radical Biology and Medicine 45, 549–561. 10.1016/j.freeradbiomed.2008.05.004.

54. Sies, H., Mailloux, R.J., and Jakob, U. (2024). Fundamentals of redox regulation in biology. Nat Rev Mol Cell Biol 25, 701–719. 10.1038/s41580-024-00730-2.

55. Veal, E.A., and Kritsiligkou, P. (2024). How are hydrogen peroxide messages relayed to affect cell signalling? Current Opinion in Chemical Biology 81, 102496. 10.1016/j.cbpa.2024.102496.

56. Stöcker, S., Van Laer, K., Mijuskovic, A., and Dick, T.P. (2018). The Conundrum of Hydrogen Peroxide Signaling and the Emerging Role of Peroxiredoxins as Redox Relay Hubs. Antioxid Redox Signal 28, 558–573. 10.1089/ars.2017.7162.

57. Zid, B.M., and O’Shea, E.K. (2014). Promoter sequences direct cytoplasmic localization and translation of mRNAs during starvation in yeast. Nature 514, 117–121. 10.1038/nature13578.

58. Winterbourn, C.C. (2008). Reconciling the chemistry and biology of reactive oxygen species. Nat Chem Biol 4, 278–286. 10.1038/nchembio.85.

59. Wood, Z.A., Poole, L.B., and Karplus, P.A. (2003). Peroxiredoxin Evolution and the Regulation of Hydrogen Peroxide Signaling. Science 300, 650–653. 10.1126/science.1080405.

60. Lim, J.M., Lee, K.S., Woo, H.A., Kang, D., and Rhee, S.G. (2015). Control of the pericentrosomal H2O2 level by peroxiredoxin I is critical for mitotic progression. J Cell Biol 210, 23–33. 10.1083/jcb.201412068.

61. Woo, H.A., Yim, S.H., Shin, D.H., Kang, D., Yu, D.-Y., and Rhee, S.G. (2010). Inactivation of Peroxiredoxin I by Phosphorylation Allows Localized H2O2 Accumulation for Cell Signaling. Cell 140, 517–528. 10.1016/j.cell.2010.01.009.

62. Cao, M., Day, A.M., Galler, M., Latimer, H.R., Byrne, D.P., Foy, T.W., Dwyer, E., Bennett, E., Palmer, J., Morgan, B.A., et al. (2023). A peroxiredoxin-P38 MAPK scaffold increases MAPK activity by MAP3K-independent mechanisms. Molecular Cell 83, 3140–3154.e7. 10.1016/j.molcel.2023.07.018.

63. Talwar, D., Messens, J., and Dick, T.P. (2020). A role for annexin A2 in scaffolding the peroxiredoxin 2–STAT3 redox relay complex. Nat Commun 11, 4512. 10.1038/s41467-020-18324-9.

64. Vo, T.N., Malo Pueyo, J., Wahni, K., Ezeriņa, D., Bolduc, J., and Messens, J. (2021). Prdx1 Interacts with ASK1 upon Exposure to H2O2 and Independently of a Scaffolding Protein. Antioxidants 10, 1060. 10.3390/antiox10071060.

65. Queiroz, R.F., Stanley, C.P., Wolhuter, K., Kong, S.M.Y., Rajivan, R., McKinnon, N., Nguyen, G.T.H., Roveri, A., Guttzeit, S., Eaton, P., et al. (2021). Hydrogen peroxide signaling via its transformation to a stereospecific alkyl hydroperoxide that escapes reductive inactivation. Nat Commun 12, 6626. 10.1038/s41467-021-26991-5.

66. Winterbourn, C.C., Peskin, A.V., Kleffmann, T., Radi, R., and Pace, P.E. (2023). Carbon dioxide/bicarbonate is required for sensitive inactivation of mammalian glyceraldehyde-3-phosphate dehydrogenase by hydrogen peroxide. Proceedings of the National Academy of Sciences 120, e2221047120. 10.1073/pnas.2221047120.

67. Tsuchiya, Y., Byrne, D.P., Burgess, S.G., Bormann, J., Baković, J., Huang, Y., Zhyvoloup, A., Yu, B.Y.K., Peak-Chew, S., Tran, T., et al. (2020). Covalent Aurora A regulation by the metabolic integrator coenzyme A. Redox Biol 28, 101318. 10.1016/j.redox.2019.101318.

68. Seisenbacher, G., Nakic, Z.R., Borràs, E., Sabidó, E., Sauer, U., de Nadal, E., and Posas, F. (2025). Redox proteomics reveal a role for peroxiredoxinylation in stress protection. Cell Rep 44, 115224. 10.1016/j.celrep.2024.115224.

69. Tiscornia, G., Singer, O., and Verma, I.M. (2006). Production and purification of lentiviral vectors. Nat Protoc 1, 241–245. 10.1038/nprot.2006.37.

70. Lasick, K.A., Jose, E., Samayoa, A.M., Shanks, L., Pond, K.W., Thorne, C.A., and Paek, A.L. (2023). FOXO Nuclear Shuttling Dynamics are Stimulus Dependent and Correspond with Cell Fate. Mol Biol Cell, mbcE22050193. 10.1091/mbc.E22-05-0193.

71. Gross, S.M., and Rotwein, P. (2015). Akt signaling dynamics in individual cells. J Cell Sci 128, 2509–2519. 10.1242/jcs.168773.

72. Livak, K.J., and Schmittgen, T.D. (2001). Analysis of relative gene expression data using real-time quantitative PCR and the 2(-Delta Delta C(T)) Method. Methods 25, 402–408. 10.1006/meth.2001.1262.

73. McQuin, C., Goodman, A., Chernyshev, V., Kamentsky, L., Cimini, B.A., Karhohs, K.W., Doan, M., Ding, L., Rafelski, S.M., Thirstrup, D., et al. (2018). CellProfiler 3.0: Next-generation image processing for biology. PLOS Biology 16, e2005970. 10.1371/journal.pbio.2005970.

74. Reyes, J., Chen, J.-Y., Stewart-Ornstein, J., Karhohs, K.W., Mock, C.S., and Lahav, G. (2018). Fluctuations in p53 Signaling Allow Escape from Cell-Cycle Arrest. Molecular Cell 71, 581–591.e5. 10.1016/j.molcel.2018.06.031.

75. Liao, Y., Smyth, G.K., and Shi, W. (2019). The R package Rsubread is easier, faster, cheaper and better for alignment and quantification of RNA sequencing reads. Nucleic Acids Res 47, e47. 10.1093/nar/gkz114.

76. Love, M.I., Huber, W., and Anders, S. (2014). Moderated estimation of fold change and dispersion for RNA-seq data with DESeq2. Genome Biol 15, 550. 10.1186/s13059-014-0550-8.

77. Xu, S., Hu, E., Cai, Y., Xie, Z., Luo, X., Zhan, L., Tang, W., Wang, Q., Liu, B., Wang, R., et al. (2024). Using clusterProfiler to characterize multiomics data. Nat Protoc 19, 3292–3320. 10.1038/s41596-024-01020-z.

