## Supplemental Figures for "A dose-dependent switch in translation capacity controls the transcription factor response to H_2_O_2_ stress"

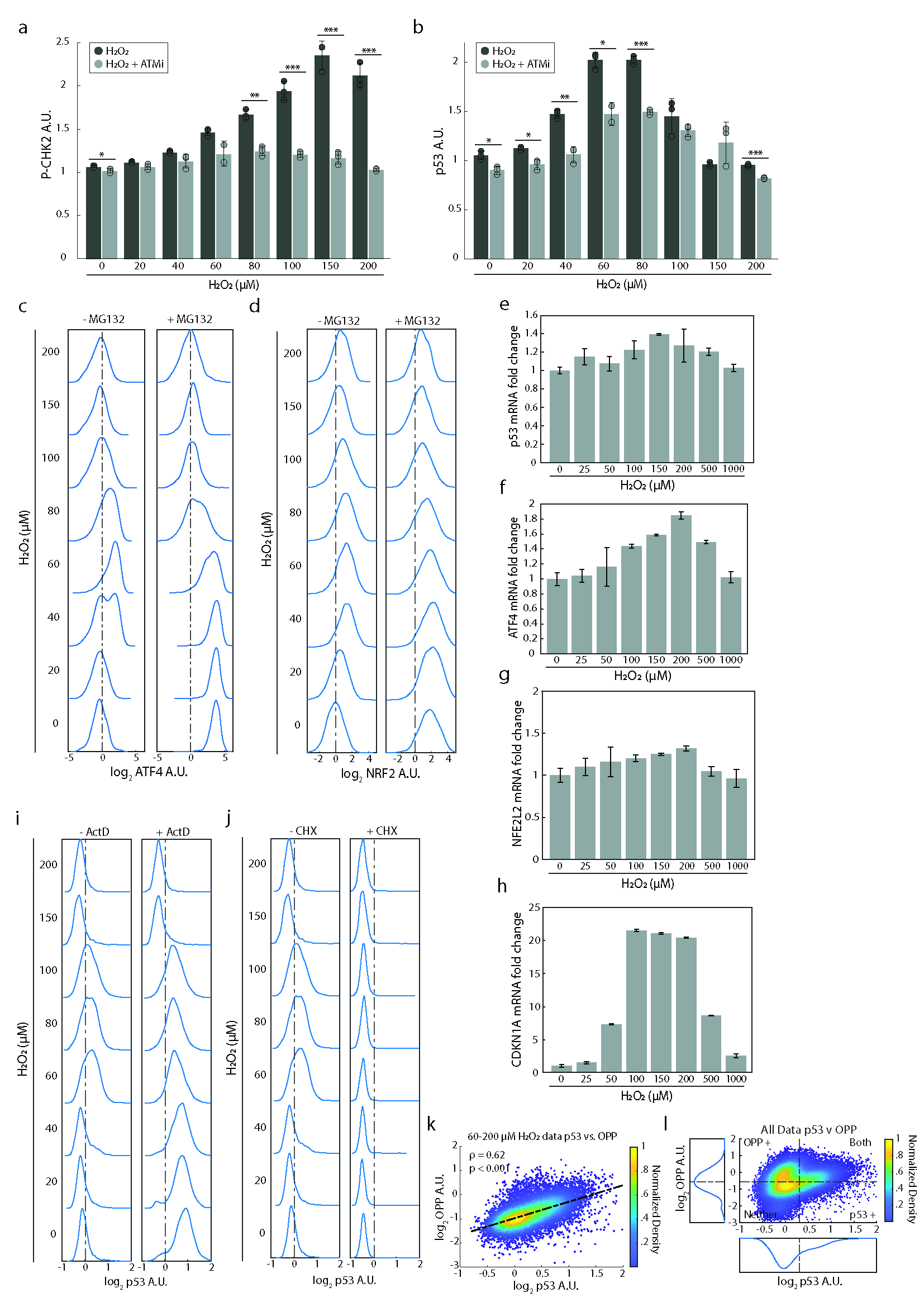
 **Supplemental Figure 1.** **(a)** Bar graph showing normalized cellular levels of P-CHK2 (T68) for MCF7 cells treated with H_2_O_2_ for 3hrs in the presence or absence of the ATM inhibitor (ATMi, KU60019, 5 µM). **(b)** Bar graph showing normalized nuclear levels of p53 for MCF7 cells treated with H_2_O_2_ in the presence or absence of ATMi (5 µM). For bar graphs (a, b), values were normalized to the H_2_O_2_-only treated condition. Error bars represent the sample standard deviation. Dots represent replicate means. P-values were acquired by first performing a two-sample t-test then adjusted for multiple hypotheses using the Benjamini-Hochberg procedure (* ≤ 0.05, ** ≤ 0.01, *** ≤ 0.001). P-Values were derived from comparing samples with and without ATMi at the same H_2_O_2_ concentration. **(c)** Population density plots of log2 nuclear ATF4 levels when treated with the indicated H_2_O_2_ concentrations for 3hrs with (+) or without (-) MG132 (5 µM). **(d)** Population density plots of log2 nuclear NRF2 levels when treated with the indicated H_2_O_2_ concentrations for 3hrs with (+) or without (-) MG132 (5 µM). **(e, f, g, h)** qPCR analysis following treatment with the indicated H_2_O_2_ concentrations for 3hrs. Graph shows fold change of the *TP53, NFE2L2*, *ATF4*, and *CDKN1A* transcripts, respectively. Error bars = sample standard deviation. **(i)** Population density plots of log2 nuclear p53 levels when treated with the indicated H_2_O_2_ concentrations for 3hrs with (+) or without (-) Actinomycin D (ActD) (1 µg/ml). **(j)** Population density plots of log2 nuclear p53 levels when treated with the indicated H_2_O_2_ concentrations for 3hrs with (+) or without (-) cycloheximide (CHX) (50 µg/ml). **(k)** Density colored scatter plot of all IF data from cells treated with 60-200 µM H_2_O_2_. The log2 nuclear p53 levels (x-axis) and cellular OPP levels (y-axis) were measured. Spearman's rank correlation was used to assess the monotonic relationship between the two variables. “ρ” = Spearman’s rank correlation coefficient. “p” = p-value from student’s t-test. **(l)** Density colored scatter plot and population density plots of IF data from all treatment conditions from the experiment in Fig. 4a. The log2 nuclear p53 levels (x-axis) and cellular OPP levels (y-axis) were measured. Activation thresholds (dashed lines) were determined using Otsu’s method. For all population density plots (c, d, i, j), values were normalized to the mean of the untreated samples (dashed line).

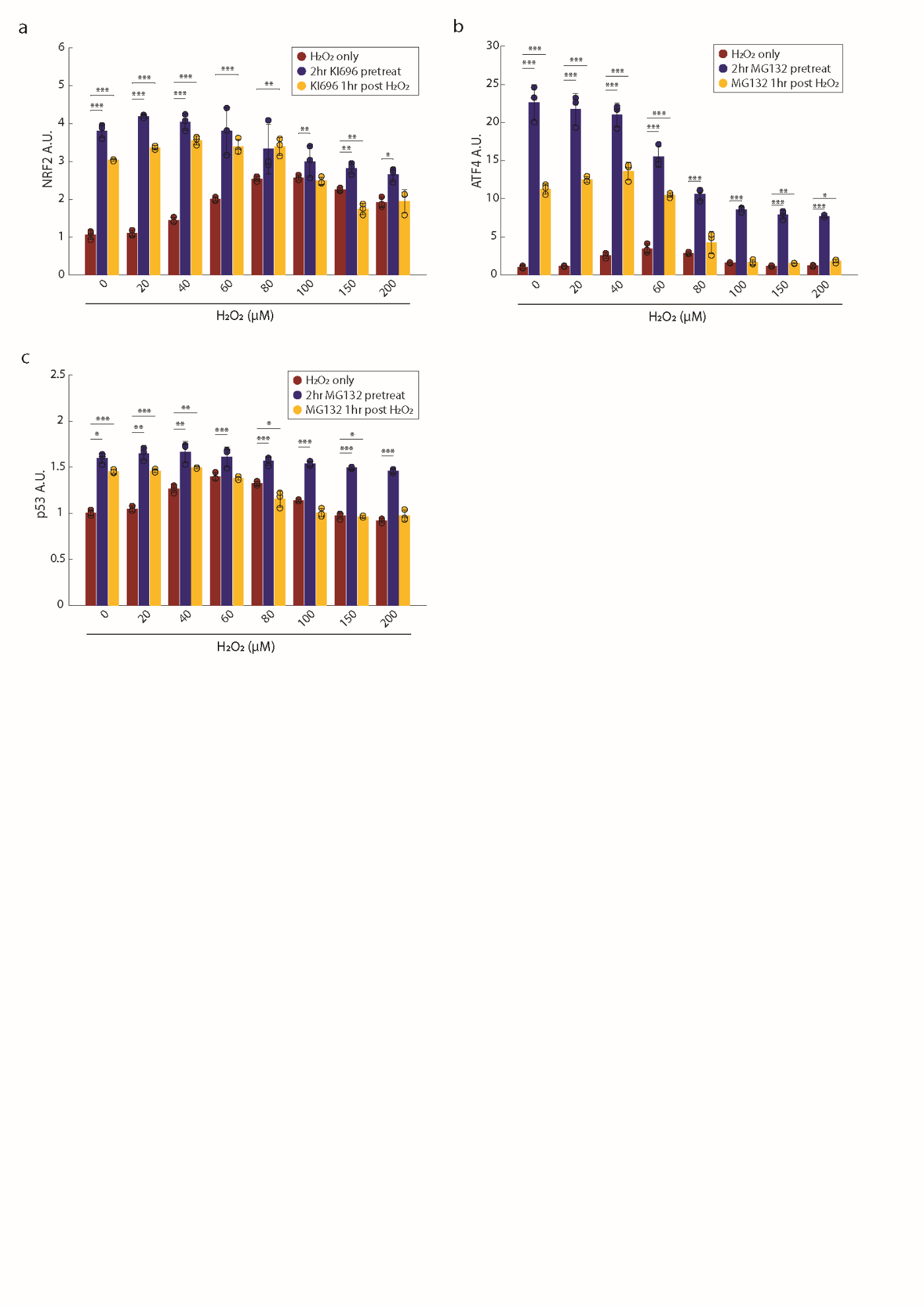
 **Supplemental Figure 2. (a)** Bar graph showing normalized nuclear NRF2 levels when treated with the indicated H_2_O_2_ concentrations alone for 3hrs or with a 2hr pretreatment with KI696 (1 µM) or treatment with KI696 1hr after H_2_O_2_. **(b)** Bar graph showing normalized nuclear ATF4 levels when treated with the indicated H_2_O_2_ concentrations alone for 3hrs or with a 2hr pretreatment with MG132 (5 µM) or treatment with MG132 1hr after H_2_O_2_. **(c)** Same as in (b) but measuring nuclear p53 levels. For bar graphs (a-c), values were normalized to the untreated condition (red bar, 0 µM). Error bars represent the sample standard deviation. Dots represent replicate means. P-values were acquired by first performing a two-sample t-test then adjusted for multiple hypotheses using the Benjamini-Hochberg procedure (* ≤ 0.05, ** ≤ 0.01, *** ≤ 0.001). P-Values were derived from comparing H_2_O_2_-only samples with the other two treatment groups at the same H_2_O_2_ concentration.

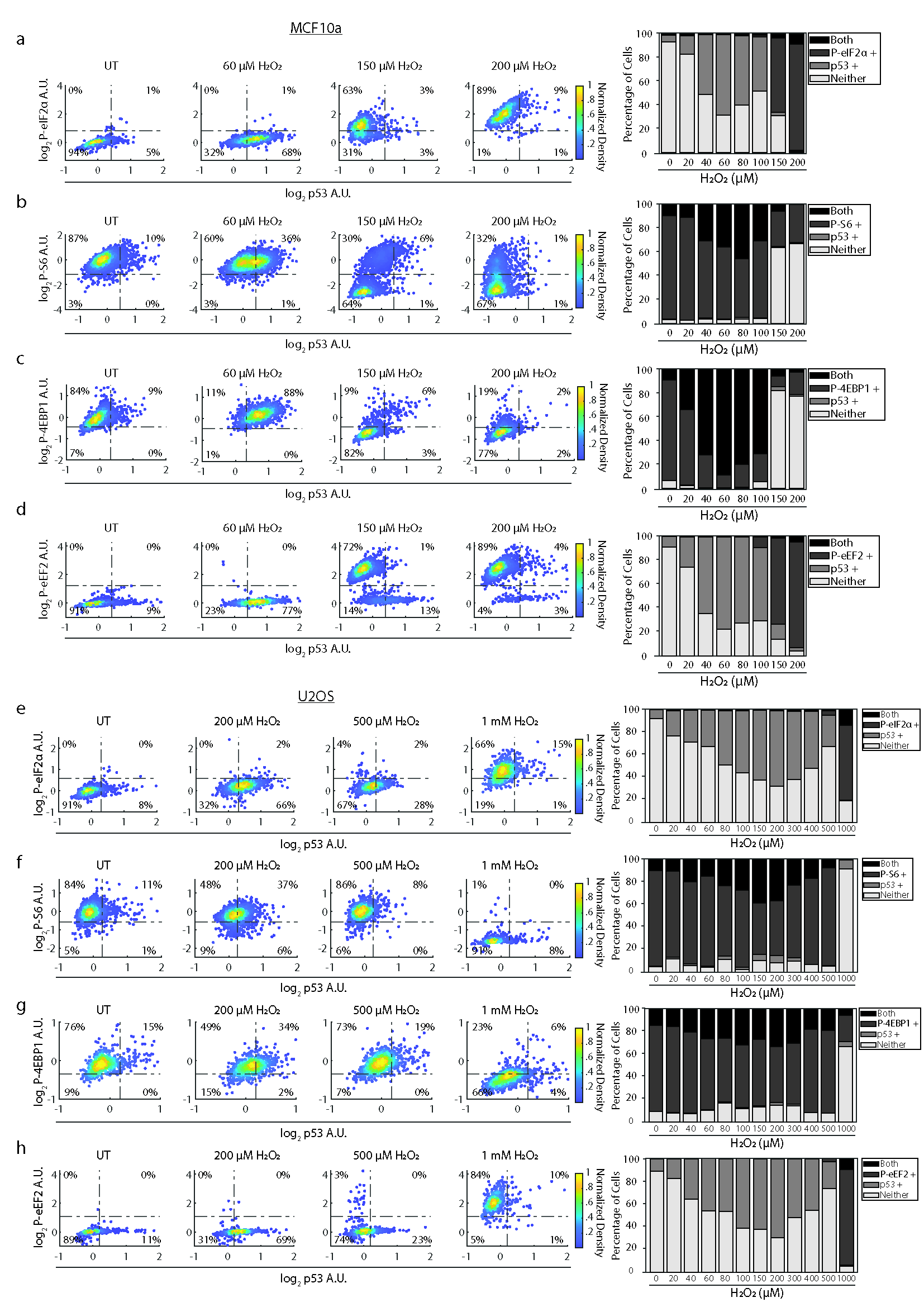
 **Supplemental Figure 3.** Density colored scatter plots of IF data from **(a-d)** MCF10A and **(e-h)** U2OS cells treated with the indicated concentrations of H_2_O_2_ for 3hrs. **(a, e)** The log2 nuclear p53 levels (x-axis) and the log2 cytoplasmic P-eIF2α (S51) (y-axis), **(b, f)** log2 nuclear p53 levels (x-axis) and the log2 cytoplasmic P-S6 (S256/236) (y-axis), **(c, g)** log2 nuclear p53 levels (x-axis) and the log2 cellular P-4EBP1 (T37/46), and **(d, h)** log2 nuclear p53 levels (x-axis) and the log2 cytoplasmic P-eEF2 (T56) (y-axis). All values were normalized to untreated controls. Activation thresholds (dashed lines) were determined using Otsu’s method on all single cell data as seen in Supp. Fig. 1k. On the right are stacked bar graphs which show the percentage of cells containing high levels of both markers (“Both”), only one (e.g. – “p53+”), or neither (“Neither”) at the indicated concentrations of H_2_O_2_.

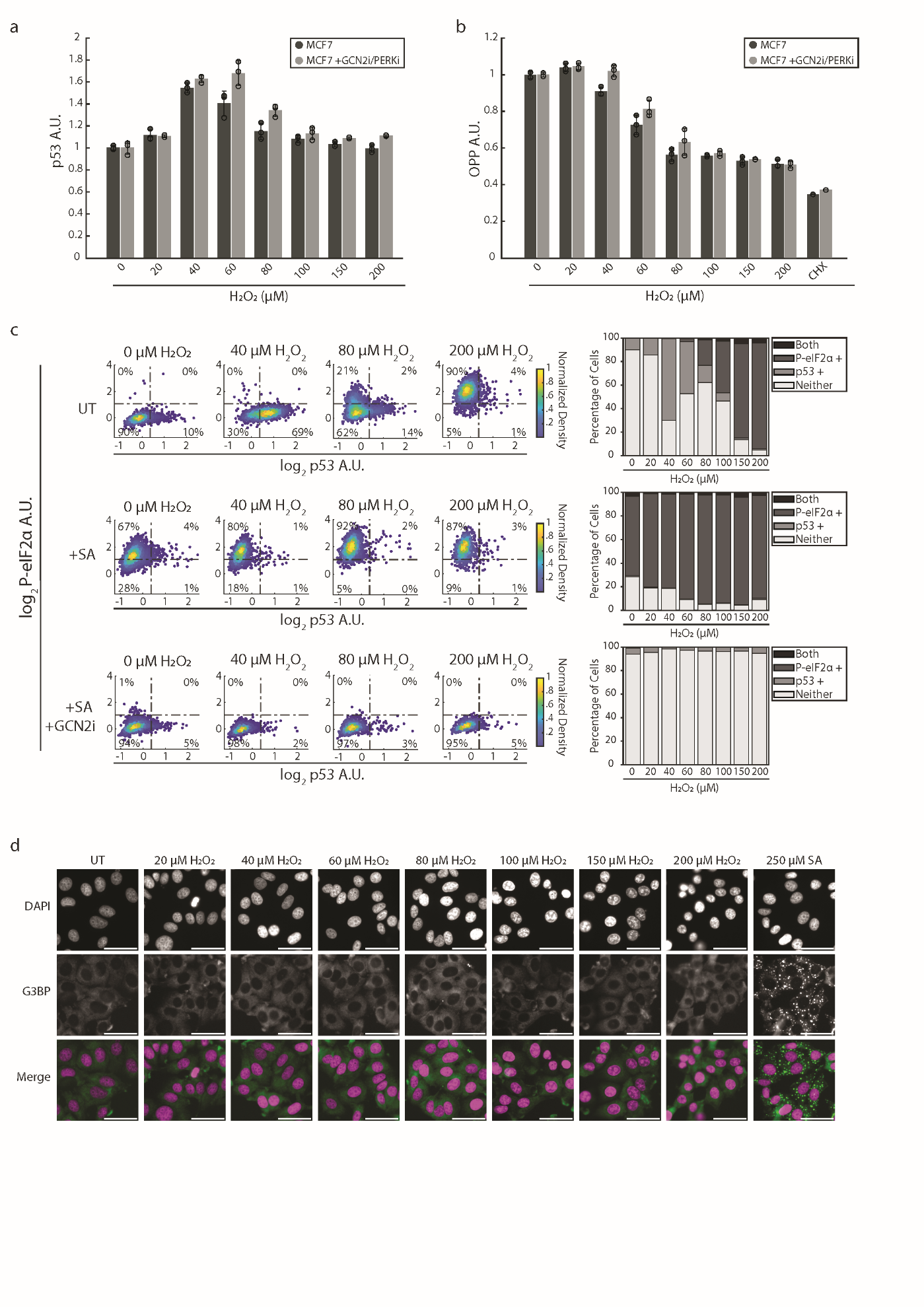
 **Supplemental Figure 4. (a)** Bar graph showing normalized nuclear levels of p53 for MCF7 cells treated with H_2_O_2_ for 3hrs in the presence or absence of PERK and GCN2 inhibitors (+PERKi 1.25 µM; +GCN2i 5 µM). **(b)** Bar graph showing normalized cellular OPP levels for MCF7 cells treated with H_2_O_2_ in the presence or absence of PERK and GCN2 inhibitors (+PERKi 1.25 µM; +GCN2i 5 µM). For bar graphs (a, b), values were normalized to the H_2_O_2_-only treated condition. Error bars represent the sample standard deviation. Dots represent replicate means. P-values were acquired by first performing a two-sample t-test then adjusted for multiple hypotheses using the Benjamini-Hochberg procedure (* ≤ 0.05, ** ≤ 0.01, *** ≤ 0.001). **(c)** MCF7 cells were treated with the GCN2 inhibitor (+GCN2i, 5 µM) for 10min before treating with 250 µM sodium arsenite (+SA) for 30min followed by H_2_O_2_ treatment for 3hrs. Density colored scatter plots of the log2 nuclear p53 levels (x-axis) and the log2 cytoplasmic P-eIF2α (S51) levels (y-axis). On the right are stacked bar graphs which show the percentage of cells containing high levels of both markers (“Both”), only one (e.g. – “p53+”), or neither (“Neither”) at the indicated concentrations of H_2_O_2_. Activation thresholds (dashed lines) were determined using Otsu’s method on all single cell data as seen in Supp. Fig. 1k. **(d)** IF staining of G3BP and DAPI counterstain after treating MCF7 cells with the indicated H_2_O_2_ concentrations for 3hrs or sodium arsenite for 30min. Scale bar = 50 µm.

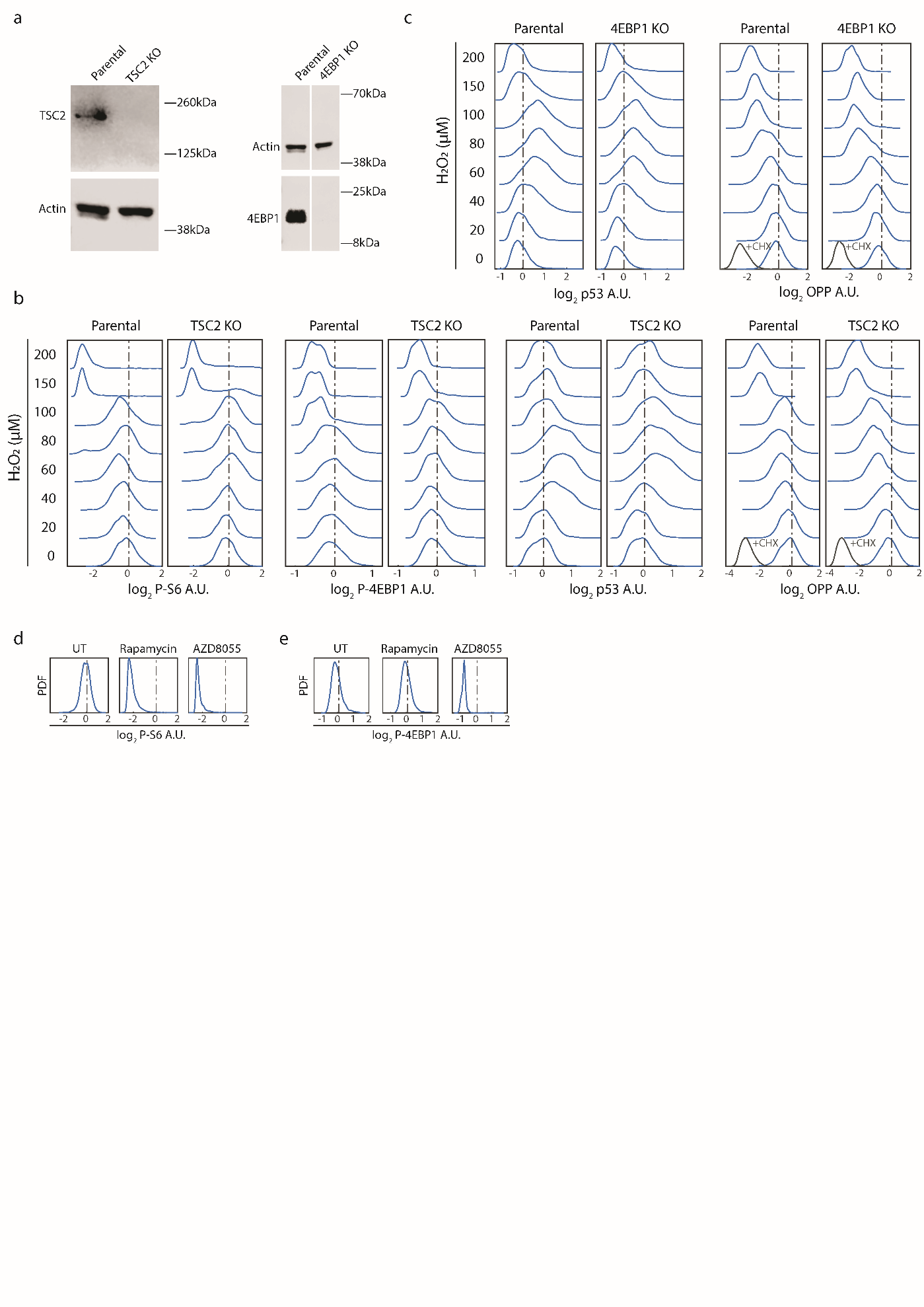
 **Supplemental Figure 5. (a)** Western blots stained for TSC2 and actin or 4EBP1 and actin in MCF7 parental cells or a clonal line containing a TSC2 or 4EBP1 CRISPR-Cas9 mediated knockout, respectively. Gap in 4EBP1 blot indicates non-adjacent lanes. **(b)** (from left to right) Population density plots of log2 cytoplasmic P-S6 (S256/236), cellular P-4EBP1 (T37/46), nuclear p53, or cellular OPP from parental or TSC2 knockout MCF7 cells treated with H_2_O_2_ for 3hrs. **(c)** Population density plots of log2 nuclear p53 (left), or cellular OPP (right) from parental or 4EBP1 knockout MCF7 cells treated with H_2_O_2_. **(d, e)** Population density plots of log2 cytoplasmic P-S6 (S256/236) and cellular P-4EBP1 (T37/46) after treating MCF7 cells with Rapamycin (220nM) or AZD8055 (100nM) for 2hrs. For all population density plots (b-e) values were normalized to the mean of the untreated samples (dashed line). +CHX = +50 µg/ml cycloheximide.

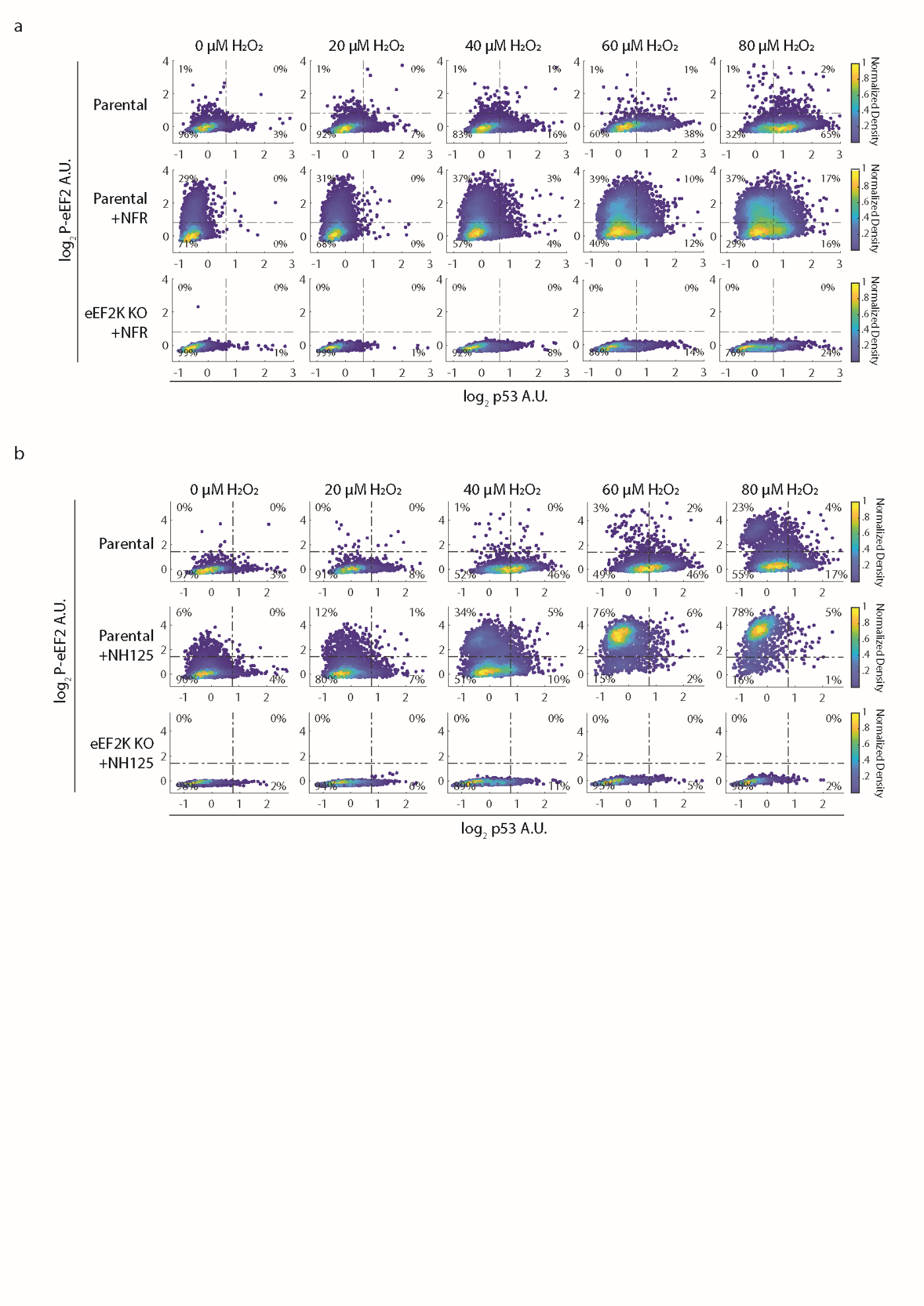
 **Supplemental Figure 6. (a)** The indicated MCF7 cell lines were treated with NFR (30 µM) for 2hrs before treating with the indicated H_2_O_2_ concentrations for 3hrs. Density colored scattered plots from IF data measuring log2 nuclear p53 levels (x-axis) and the log2 cytoplasmic P-eEF2 (T56) (y-axis). **(b)** As in (a), but NH125 (5uM) instead of NFR. Activation thresholds (dashed lines) were determined using Otsu’s method on all single cell data as seen in Supp. Fig. 1k.

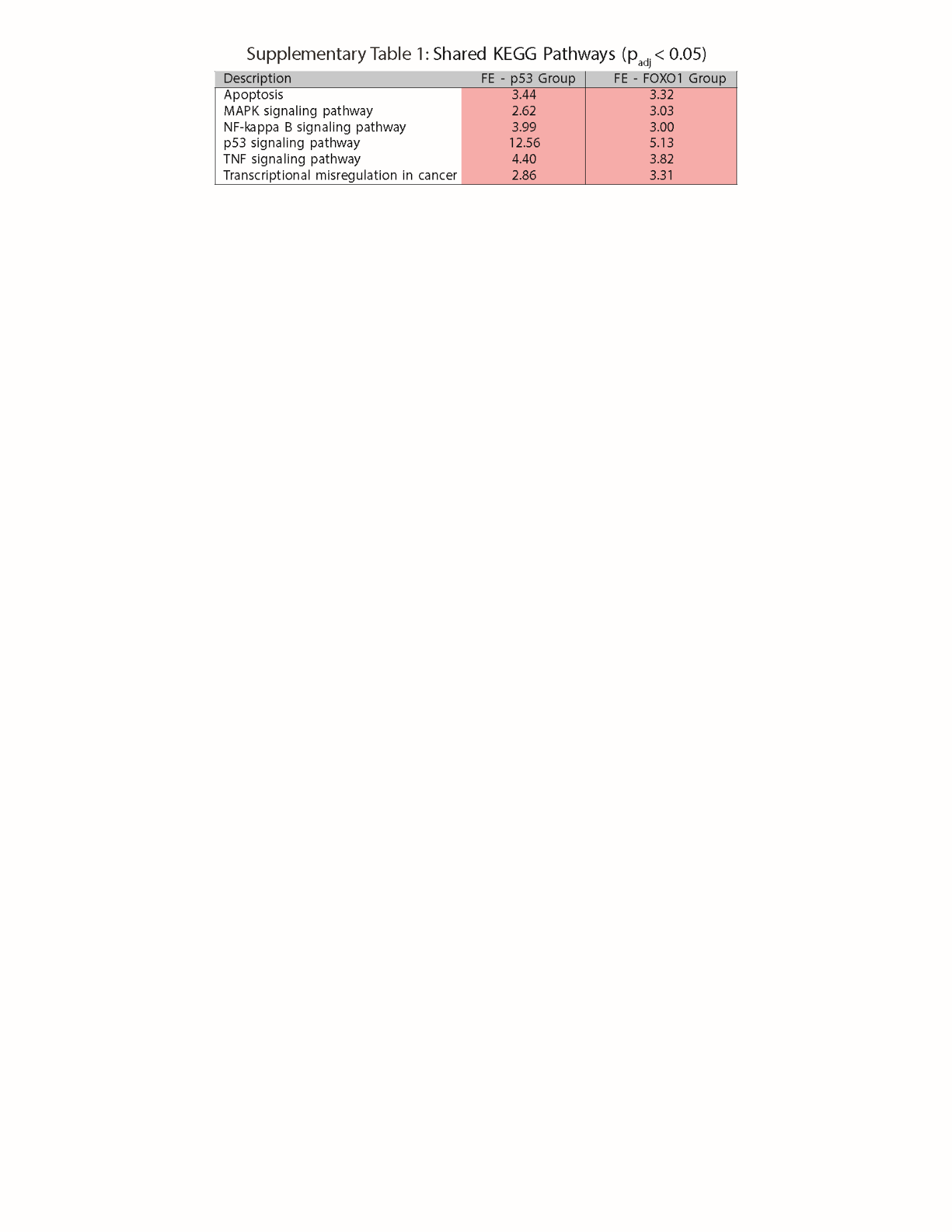
 **Supplemental Table 1.** KEGG analysis using significantly enriched genes (p_adj_ < 0.01, |Log2FoldChange| > 1) was used to find overlapping pathways enriched in cells with either p53 active (“p53 Group”) or suppressed (“FOXO1 Group”) after H_2_O_2_ treatment. “FE” = Fold Enrichment. Red color indicates a positive enrichment.

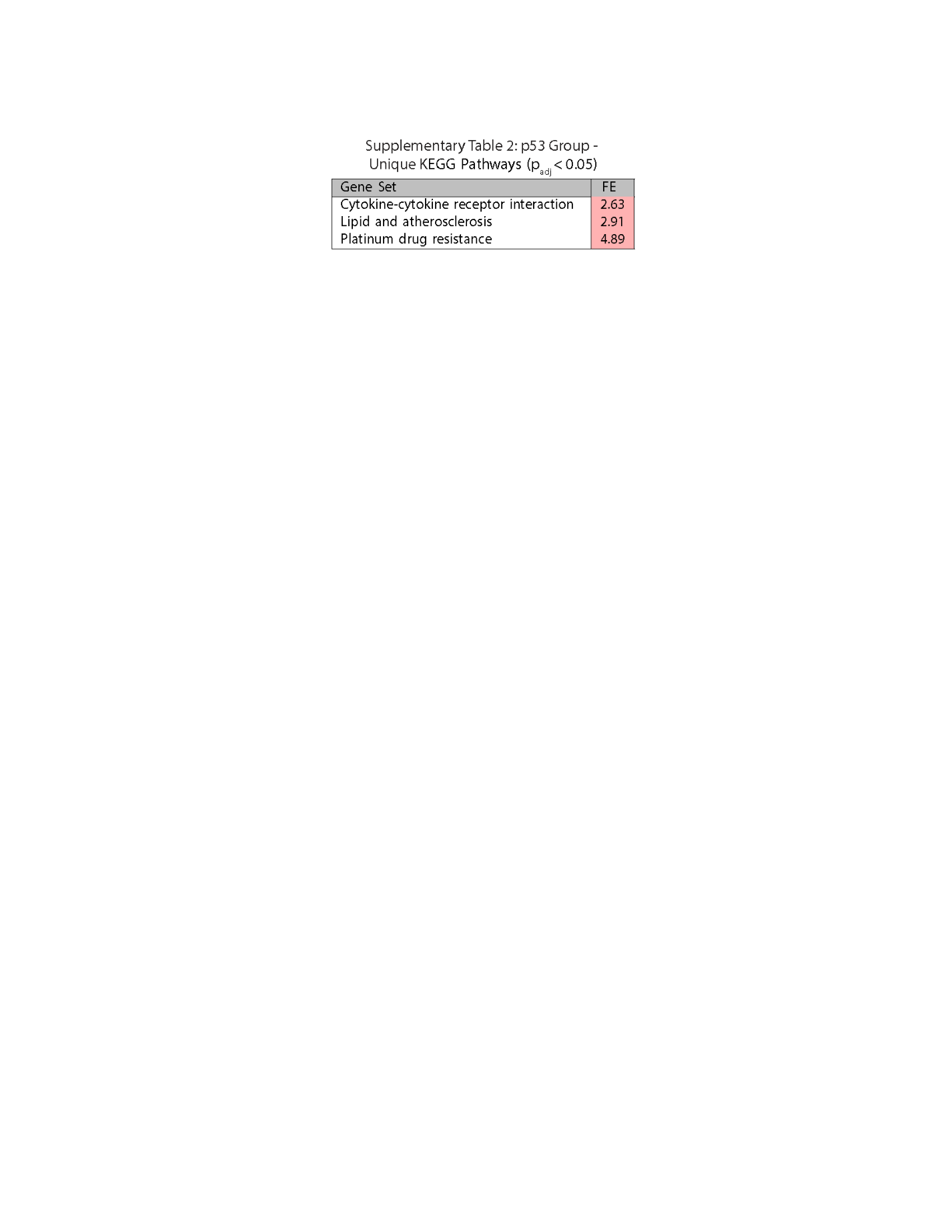
 **Supplemental Table 2.** KEGG analysis using significantly enriched genes (p_adj_ < 0.01, |Log2FoldChange| > 1) was used to find unique pathways enriched in cells with p53 active (“p53 Group”) after H_2_O_2_ treatment. “FE” = Fold Enrichment. Red color indicates a positive enrichment.

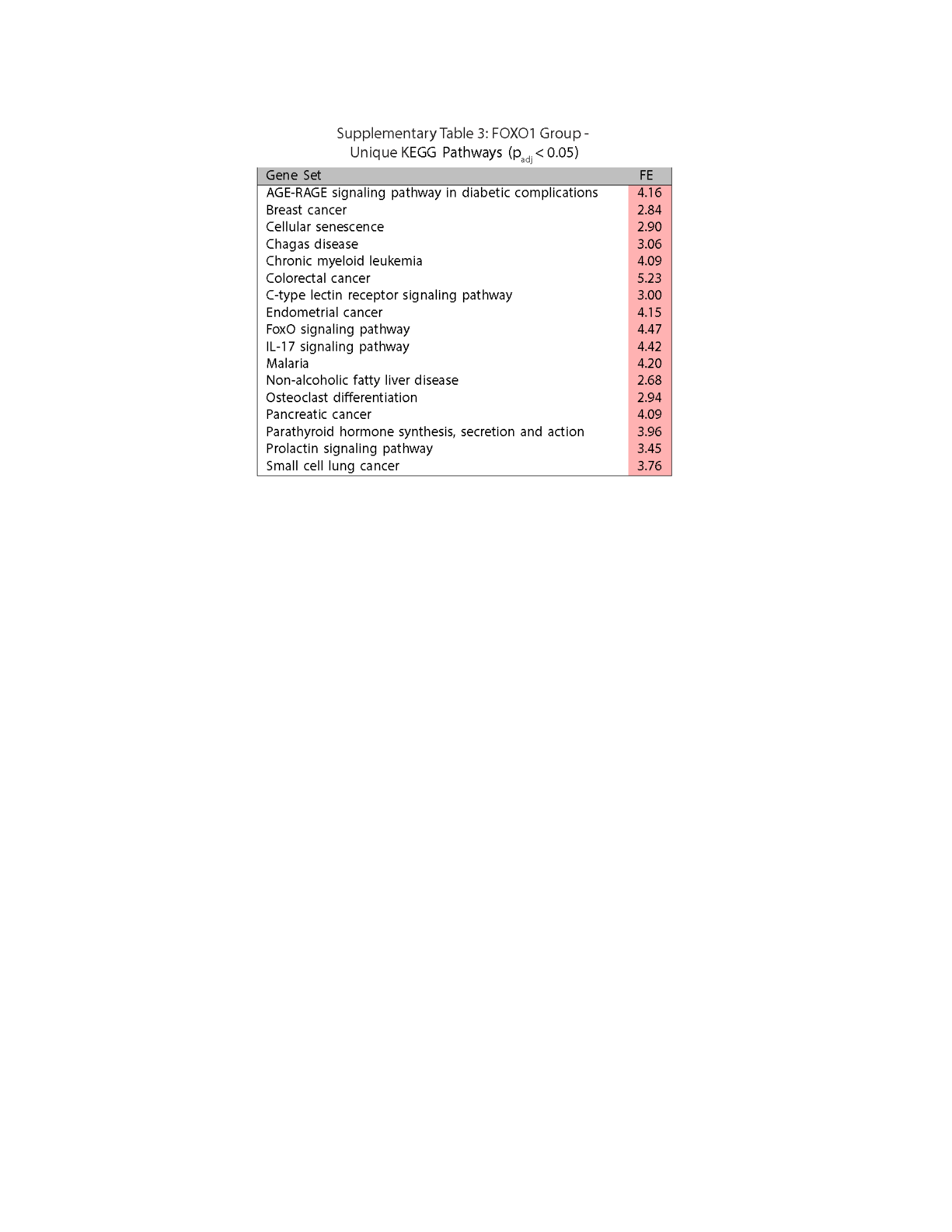
 **Supplemental Table 3.** KEGG analysis using significantly enriched genes (p_adj_ < 0.01, |Log2FoldChange| > 1) was used to find unique pathways enriched in cells with FOXO1 active (“FOXO1 Group”) after H_2_O_2_ treatment. “FE” = Fold Enrichment. Red color indicates a positive enrichment.

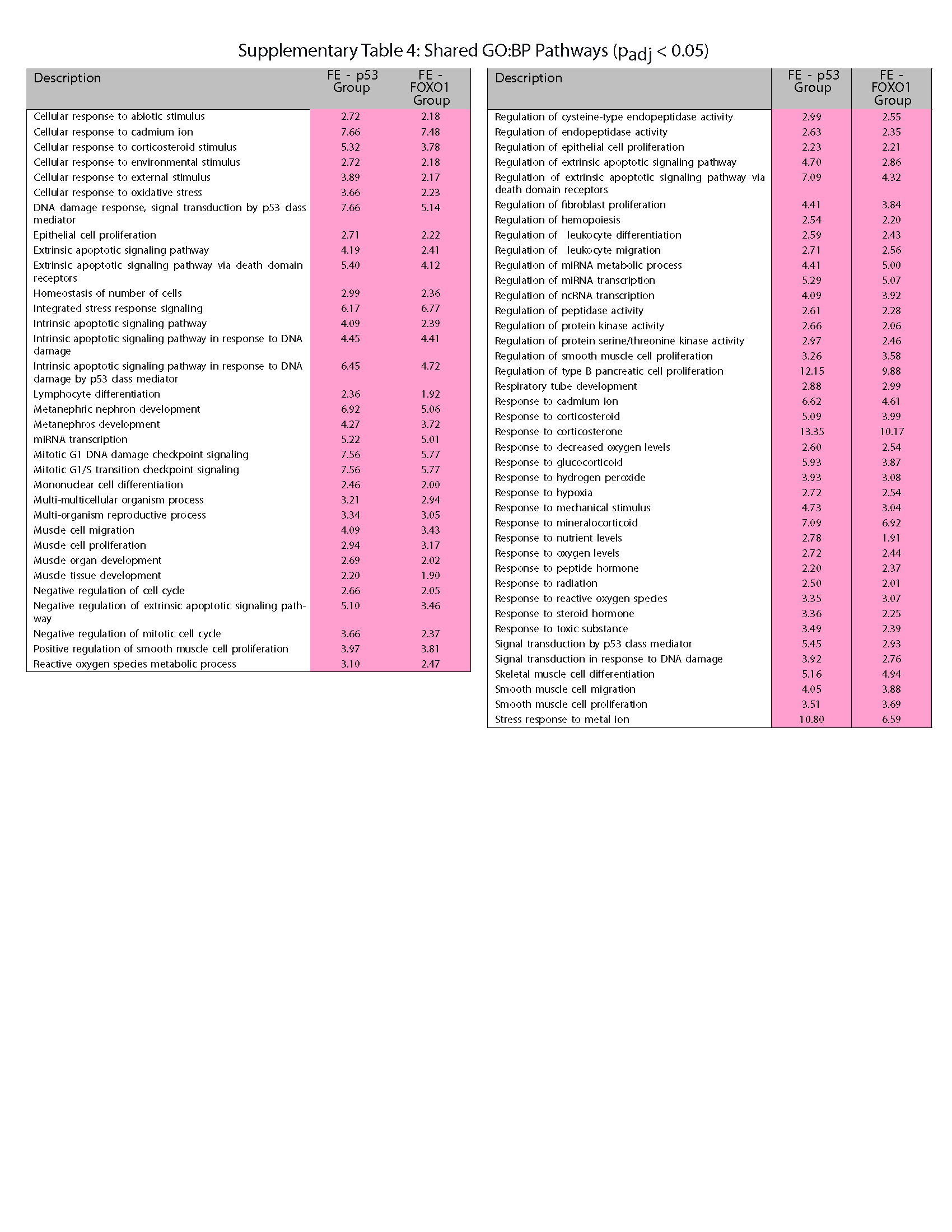
 **Supplemental Table 4.** Gene Ontology analysis using significantly enriched genes (p_adj_ < 0.01, |Log2FoldChange| > 1) and the Biological Processes (“GO:BP”) category was used to find overlapping pathways enriched in cells with either p53 active (“p53 Group”) or suppressed (“FOXO1 Group”) after H_2_O_2_ treatment. “FE” = Fold Enrichment. Red color indicates a positive enrichment.

(Attached data files)

**Supplementary Data 1.** Gene Ontology analysis using significantly enriched genes (p_adj_ < 0.01, |Log2FoldChange| > 1) and the Biological Processes (“GO:BP”) category was used to find unique pathways enriched in cells with p53 active (“p53 Group”) after H_2_O_2_ treatment. (“SuppData1 - p53 Group GOBP Pathways.csv”)

**Supplementary Data 2.** Gene Ontology analysis using significantly enriched genes (p_adj_ < 0.01, |Log2FoldChange| > 1) and the Biological Processes (“GO:BP”) category was used to find unique pathways enriched in cells with FOXO1 active (“FOXO1 Group”) after H_2_O_2_ treatment. (“SuppData2 - FOXO1 Group GOBP Pathways.csv”)
